# (p)ppGpp buffers cell division when membrane fluidity decreases in *Escherichia coli*

**DOI:** 10.1101/2024.03.27.586906

**Authors:** Vani Singh, Rajendran Harinarayanan

**Affiliations:** Center for DNA Fingerprinting and Diagnostics, Hyderabad, India 500039; Manipal Academy of Higher Education, Manipal, India 576104

## Abstract

Maintenance of fluidity an inherent property of biological membranes by homeoviscous adaptation is important for optimal functioning of membrane-associated processes. Homeoviscous adaptation in *E. coli* involves an increase in the concentration of unsaturated fatty acid, cis-vaccenic acid (18:1) with decrease in temperature and *vice versa*. Lowering unsaturated fatty acid synthesis by inactivation of FadR reduced the proportion of unsaturated fatty acids in the membrane. In this study we show that when the proportion of unsaturated fatty acids in the membrane was reduced, cell division was dependent on the guanine nucleotide analogous (p)ppGpp. Combined expression of cell division genes *ftsQ*, *ftsA* and *ftsZ* from plasmid rescued the growth defect that was associated with cell filamentation followed by lysis. To our knowledge, this is the first report of (p)ppGpp mediated regulation needed for the adaptation to membrane fluidity loss in bacteria.

## Introduction

For bacterial cells, nutritional content of growth medium is an important determinant of macromolecular synthesis capacity and growth rate (1). In *E. coli* and other bacteria, the guanine nucleotide analogs, pppGpp (guanosine penta phosphate) and ppGpp (guanosine tetra phosphate), collectively referred as (p)ppGpp are synthesized by the widely conserved RSH protein family (2) and elicit stringent response to variety of nutritional stresses (3). In *E. coli*, (p)ppGpp concentration correlates inversely with growth rate even in the absence of stress (4) and has been proposed as a primarily negative regulator of growth rate (5). In support of this, there is evidence for negative regulation of macromolecular synthesis processes, namely, replication, transcription and translation directly or indirectly by (p)ppGpp (3, 6).

Cell size correlates positively with growth rate and is determined by the balance of biomass synthesis rate and rate of cell cycle progression, which is the volume added between one cell division to the next. Under steady state growth conditions, there is constant volume addition to cells independent of their size at birth (7, 8). Although mechanisms are not understood, studies have implicated (p)ppGpp in the regulation of cell division (9, 10). Regulation of growth rate and cell division by (p)ppGpp would make it an ideal candidate for cell size regulation. By artificially regulating (p)ppGpp level, a recent study showed its effect on cell size precedes its effect on growth rate, that is, the volume added during cell cycle progression correlates with (p)ppGpp level rather than growth rate (11). Cells of Δ*relA* Δ*spoT* mutant that lack (p)ppGpp, continued to show cell size increase with growth rate increase, but were larger than isogenic wild type cells in rich medium (12). This suggested even low basal levels of (p)ppGpp played a role in cell size regulation.

Recent studies have highlighted the importance of membrane biogenesis in cell size determination – an increase or decrease in fatty acid biosynthesis, increased and decreased respectively the cell size (12, 13). (p)ppGpp accumulates in response to fatty acid starvation (14, 15) and negatively regulates fatty acid and phospholipid biosynthesis (16, 17). Evidence for (p)ppGpp being the linchpin co-ordinating macromolecular biosynthesis with membrane biogenesis in *E. coli* was presented in Vadia *et. al*., (12). Co-ordination of lipid synthesis and cell growth by stringent response was reported in *B. subtilis* (18) and could be a widely conserved across bacteria.

FadR is a transcription factor central to fatty acid metabolism. It represses the expression of *fad* genes involved in fatty acid degradation and activates expression of *fabA* and *fabB* genes involved in unsaturated fatty acid biosynthesis (19). Interaction of acyl-CoA to FadR regulates the DNA binding of FadR (20). FadR activates the expression of *fabHDG* genes encoding the enzymes that carry out initiation and elongation of fatty acid biosynthesis (17). FadR overproduction increased the expression of genes involved in fatty acid biosynthesis (21) and enhanced fatty acid production (22).

Over-expression of FadR overcame the cell size defect associated with increase in (p)ppGpp, indicating cell size regulation through membrane biogenesis occurred downstream of (p)ppGpp regulation and possibly uncoordinated from (p)ppGpp regulated metabolic processes (12). If membrane biogenesis was the primary determinant of cell size, independent of (p)ppGpp, then inhibition of fatty acid biosynthesis may be expected to decrease the cell size of (p)ppGpp deficient strain. However, inhibition of fatty acid biosynthesis with cerulenin led to the loss of membrane integrity and cell death (12).

In this study, we systematically examined the consequences of inhibiting fatty acid biosynthesis and found that the (p)ppGpp deficient strains were sensitive to perturbations that decrease the proportion of unsaturated fatty acid content in the membrane. The growth defect of (p)ppGpp deficient strain was due to the inability to sustain cell division when the unsaturated fatty acid content of the membrane was reduced. Combined expression of *ftsQAZ* genes from plasmid rescued the cell division defect, suggesting positive regulation of cell division by (p)ppGpp was required for the divisome to function under low fluidity conditions.

## RESULTS

### Reducing (p)ppGpp content impaired the growth of Δ*fadR* mutant

Since FadR plays a central role in fatty acid metabolism, we used the Δ*fadR* to ask if changes in (p)ppGpp concentration affected the growth of this strain. The plating efficiency of Δ*relA* Δ*fadR* double mutant was compared with that of Δ*relA* and Δ*fadR* single mutants in LB medium. While the single mutant strains grow well at 25°C, 30°C, and 37°C, the Δ*relA* Δ*fadR* strain showed a clear decrease in plating efficiency at 25°C, and gradually recovered with increase in growth temperature (Fig. 1A). The isogenic wild type strain MG1655, did not show growth defect at any temperature (data not shown).

**Figure 1.**
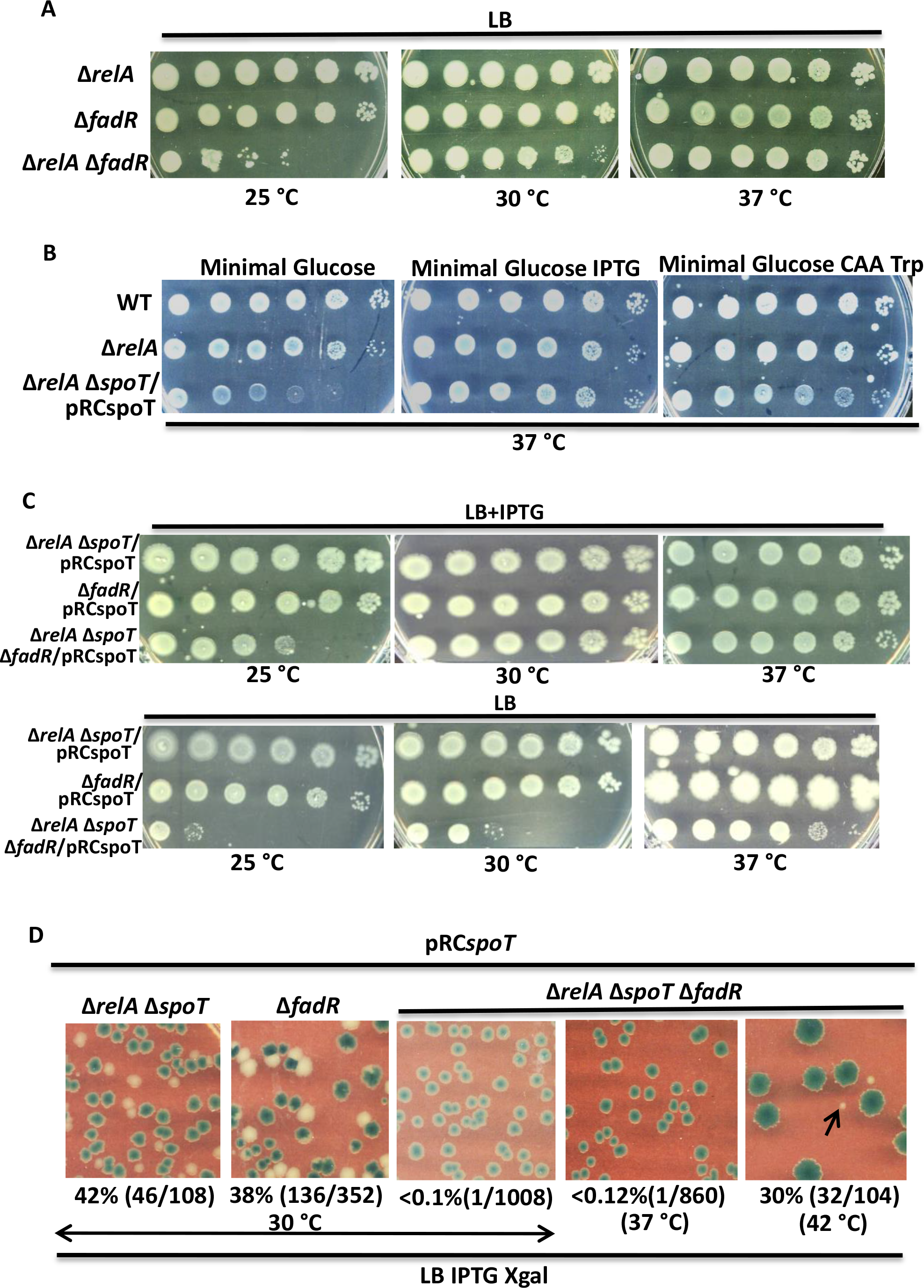
Genetic changes that reduce (p)ppGpp pool impaired the growth of Δ*fadR* mutant at low temperature. Strains whose relevant genotypes are mentioned were cultured to stationary phase in LB medium or LB medium containing IPTG at 37⁰C, washed and serially diluted with minimal A medium and spotted on LB agar plates (A, C), minimal A agar plates (B), suitable dilution was spread on LB medium containing IPTG (1mM) and X-Gal (50µg/ml) to estimate frequency of pRCspoT plasmid loss by quantifying the number of blue/white colonies, representative panels are shown (D). The percentage of white colonies and the number of white colonies out of total colonies counted (blue+white) are shown below each panel. The incubation temperature and supplements added to growth medium are mentioned besides the panels. Final concentration of CAA (casamino acids) and Trp (tryptophan) was 0.2% and 40µg/ml respectively. The strains are MG1655 (WT, VS2), Δ*relA* (VS3), Δ*fadR* (VS11), Δ*relA* Δ*fadR* (VS33), Δ*relA* Δ*spoT* /pRC*spoT* (AN120), Δ*fadR* /pRC*spoT* (VS9), Δ*relA* Δ*spoT* Δ*fadR*/pRC*spoT* (VS6). All strains have the Δ*lacZYAI*::FRT allele.

We studied the effect of Δ*fadR* mutation in the Δ*relA* Δ*spoT*/pRC*spoT* strain, where (p)ppGpp concentration can be modulated through IPTG regulated *spoT* expression from pRC*spoT* plasmid (23). In this construct made using plasmid pRC7 (24), an unstable single-copy vector derived from F plasmid, the *spoT* and *lacZ* genes are expressed from the IPTG-regulated *lac* promoter. An IPTG dependent increase in the (p)ppGpp content in Δ*relA* Δ*spoT* /pRC*spoT* strain can be deduced from an IPTG-dependent rescue of multiple amino acid auxotrophy of Δ*relA* Δ*spoT* strain (Fig 1B) (25). The poor growth of Δ*relA* Δ*spoT*/pRC*spoT* strain, as compared to Δ*relA* strain in minimal glucose medium lacking IPTG, reflected the lower basal (p)ppGpp pool in the former strain in the absence of *spoT* induction. To study growth of Δ*fadR* mutant under (p)ppGpp depleted condition, the Δ*relA* Δ*spoT* Δ*fadR* /pRC*spoT* strain was constructed and its plating efficiency was examined relative to isogenic *fadR*^+^ (Δ*relA* Δ*spoT* /pRC*spoT*) and *relA*^+^ *spoT*^+^ (Δ*fadR* Δ*lac*/pRC*spoT*) strains at different growth temperatures (Fig. 1C). The latter two strains did not exhibit growth defect at all tested temperatures and this was independent of *spoT* expression. On the other hand, the Δ*relA* Δ*spoT* Δ*fadR* /pRC*spoT* strain showed plating defect at 25°C in the presence of IPTG and both 25°C and 30°C, in the absence of IPTG. These results showed, in the absence of FadR function, lowering the (p)ppGpp concentration impaired growth at 30°C and lower.

To study growth of Δ*fadR* mutant in the absence of (p)ppGpp, we took advantage of the unstable nature of pRC*spoT* plasmid and studied its segregation. The plasmid segregation assay is based on the rationale that an essentiality of plasmid encoded function can stabilize a normally unstable plasmid. During unselected growth, random plasmid segregation results in emergence of cells retaining or losing the plasmid (see methods for detailed description). When such cultures are diluted and spread on plates, cells lacking the plasmid grow to form white colonies, cells with plasmid form blue colonies or sectored colonies if plasmid segregation occurs during the growth on plate (it should be noted the strains have the Δ*lacZYAI*::FRT deletion on the chromosome). Strains cultured overnight without selection for pRCspoT, were diluted and spread on plates. Following incubation at 30°C, loss of pRCspoT plasmid was evident in the Δ*relA* Δ*spoT* genetic background (Δ*relA* Δ*spoT*/pRC*spoT* strain) and Δ*fadR* genetic background (Δ*fadR* /pRC*spoT* strain) after 36 hrs of incubation, but this was not the case in the Δ*relA* Δ*spoT* Δ*fadR* genetic background (Δ*relA* Δ*spoT* Δ*fadR*/pRC*spoT* strain) even up to 72 hrs of incubation (Fig. 1D). Plasmid loss was observed in the latter strain after 36 hr of incubation at 42°C but not at 37°C (Fig. 1D). The Δ*relA* Δ*spoT* Δ*fadR* strain (white colonies) was slow growing at 42°C relative to those expressing spoT (blue colonies).

To sum up, the growth of Δ*fadR* mutant showed dependence on (p)ppGpp and temperature. (i) it grew at 30⁰C but not 25⁰C with SpoT as the sole source of (p)ppGpp (ii) when more severely depleted for (p)ppGpp it grew at 37 ⁰C but not 30 ⁰C; (iii) in the absence of (p)ppGpp it grew at 42⁰C but not 37⁰C. It can be said, growth of Δ*fadR* mutant below 42⁰C was dependent on (p)ppGpp - lower the growth temperature, greater the (p)ppGpp requirement for growth.

### Genetic changes that increase unsaturated fatty acid synthesis rescued the growth defect of (p)ppGpp deficient Δ*fadR* strains

An *E. coli* genomic library constructed in the multi-copy plasmid pACYC184 was introduced into Δ*relA* Δ*spoT* Δ*fadR* /pRCspoT strain to identify gene over-expression suppressors of growth defect. Two plasmid clones that supported IPTG independent growth at 30°C were identified. Sequencing the plasmid-chromosomal DNA junction revealed the presence of ‵*relA- mazE-mazF*′ contig. This plasmid rescued the multiple amino acid auxotrophy of Δ*relA* Δ*spoT* strain (data not shown), indicating the truncated *relA* gene (lacking the first 9 codons) was capable of (p)ppGpp synthesis and therefore could rescue the growth defect. This result reinforced the idea that insufficient (p)ppGpp was responsible for growth defect of Δ*relA* Δ*spoT* Δ*fadR*/pRCspoT strain at 30°C. The second plasmid carried the contig, *cspH*′*-cspG*- *ymcF*-*ymcE*-*gnsA-yccM-*‵*torS*. Amongst the annotated genes was *gnsA,* the over-expression of which has been reported to increase unsaturated fatty acid synthesis (26, 27). *gnsB*, a sequence homolog of *gnsA* can also increase unsaturated fatty acid (27). We hypothesized, decrease in unsaturated fatty acid content could be the primary cause for growth defect of (p)ppGpp deficient Δ*fadR* mutants at low temperature. This was tested using the *gnsA* and *gnsB* plasmid clones from the ASKA library (28). Both plasmids, pCAgnsA and pCAgnsB, but not the vector rescued the growth defect of Δ*relA* Δ*fadR* strain at 25°C (Fig. 2A). The plasmids supported growth of Δ*relA* Δ*spoT* Δ*fadR* strain at 30°C as seen from segregation of white colonies in the *ΔrelA ΔspoT* Δ*fadR*/pRCspoT strain in the presence of pCA*gnsA* or pCA*gnsB*, but not the vector pCA24N (Fig. 2B, panels i to iii).

**Figure 2.**
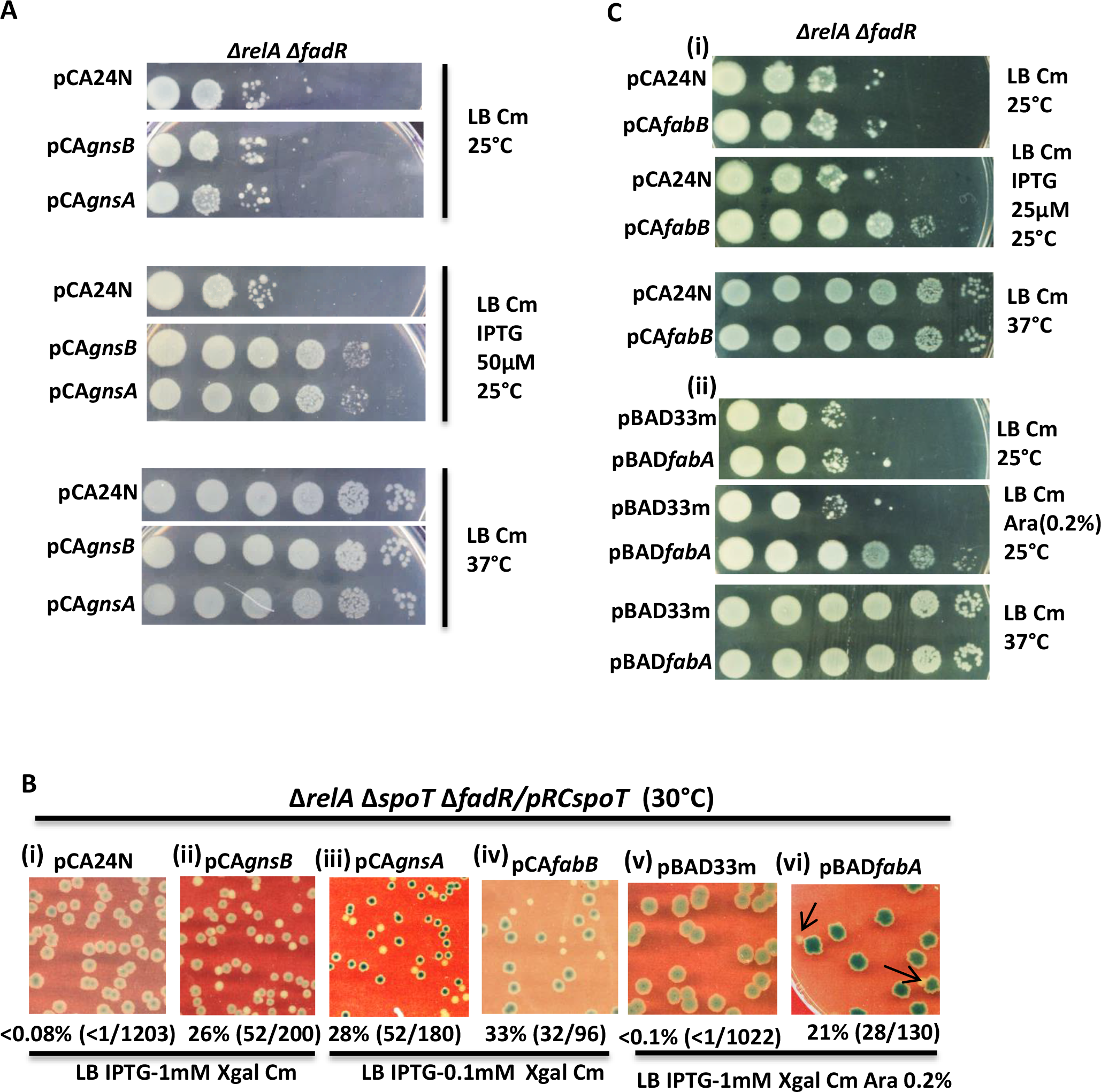
Genetic changes that increase unsaturated fatty acid biosynthesis rescued the growth defect of (p)ppGpp deficient Δ*fadR* strains at low temperature. Strains whose relevant genotypes are mentioned were cultured to stationary phase in LB medium containing appropriate antibiotics and IPTG at 37⁰C, but without selection for pRCspoT, washed and serially diluted with minimal A and spotted on LB agar plates (A & C). Suitable dilutions were spread on LB agar plates containing IPTG and X-Gal to estimate the frequency of pRCspoT plasmid loss by quantification of blue/white colonies (B). The incubation temperature, inducers used for expression of plasmid borne genes and antibotic used for plasmid selection are mentioned besides the panel. The percentage of white colonies and the number of white colonies out of total colonies counted (blue+white) are shown below each panel. The strains used are, Δ*relA* Δ*fadR*/pCA24N (VS149), Δ*relA* Δ*fadR*/pCA*gnsB* (VS134), Δ*relA* Δ*fadR*/ pCA*gnsA* (VS138), Δ*relA* Δ*fadR*/pCA*fabB* (VS136), Δ*relA* Δ*fadR*/pBAD33m (VS135), Δ*relA* Δ*fadR*/pBAD*fabA* (VS137), Δ*relA* Δ*spoT ΔfadR* /pRC*spoT*/pCA24N (VS15), Δ*relA* Δ*spoT ΔfadR* / pRC*spoT*/pCA*gnsB* (VS14), Δ*relA* Δ*spoT* Δ*fadR* /pRC*spoT* / pCA*gnsA* (VS22), Δ*relA* Δ*spoT* Δ*fadR* / pRC*spoT*/pCAfabB (VS23), Δ*relA* Δ*spoT* Δ*fadR*/pRC*spoT*/pBAD33m (VS120), Δ*relA* Δ*spoT* Δ*fadR* / pRC*spoT*/pBAD*fabA* (VS119). All strains have the Δ*lacZYAI*::FRT allele.

The *fabA* and *fabB* genes are necessary for unsaturated fatty acid biosynthesis, and their expression is reduced in the *fadR* mutant (29, 30). Using the *fabA* or *fabB* clone from the ASKA collection, we tested if expression of each gene, rescued the growth defect of (p)ppGpp deficient Δ*fadR* strains. The *fabB* clone rescued growth defect of Δ*relA* Δ*fadR* strains at 25°C (Fig. 2C panel i) after IPTG induction and the Δ*relA* Δ*spoT* Δ*fadR* strain at 30°C (Fig. 2B panel iv). The *fabA* clone did not rescue the growth defect, but also failed to rescue the temperature sensitivity of *fabA*(ts) strain at 42°C (data not shown). The *fabA* gene was likely to be non-functional. When *fabA* gene PCR amplified from MG1655, was cloned in the pBAD24 vector and tested, arabinose dependent rescue of growth defect was observed (Fig. 2B panels v & vi, panels in Fig. 2C ii). This plasmid rescued the temperature sensitivity of *fabA*(ts) strain in the presence of arabinose (Fig. S1).

To find out if there was an increase in the unsaturated fatty acid content associated with growth rescue, we examined fatty acid composition by fatty acid methyl ester (FAME) analysis (Table 1). Consistent with an earlier report (31), the proportion of unsaturated and saturated fatty acids decreased and increased respectively, in the Δ*fadR* mutant (compare rows 1 and 2). The proportion of fatty acid species in the Δ*relA* Δ*spoT* strain was largely similar to wild type (rows 1 and 3), a slight decrease in 18:1 (cis-vaccenic acid) was notable. In (p)ppGpp depleted and growth arrested Δ*relA* Δ*spoT* Δ*fadR*/pRCspoT strain (following growth of the strain without IPTG, see methods) the proportion of fatty acid species were similar to that in Δ*fadR* strain (rows 2 to 4). This indicated, (p)ppGpp depletion did not alter the relative proportion of unsaturated versus saturated fatty acid in the presence or absence of FadR. Expression of *gnsA* in the Δ*relA* Δ*spoT* Δ*fadR* strain restored the proportion of 16:1 and 18:1 to that seen in wild type and lowered the proportion of 16:0 (compare rows 1, 5 and 6). *fabA* or *fabB* expression primarily increased 16:1 and to a smaller extent 18:1and decreased the proportion of 16:0 (compare rows 5, 7 to 9).

**Table 1.** Profile of major fatty acids in strains grown in LB medium at 30°C.

| Strains |  | Major Fatty Acid Species |  |  |  |  |  |
| --- | --- | --- | --- | --- | --- | --- | --- |
| | | 16:1 | 18:1 | 16:0 | 14:0 | 12:0 | $\frac{(16:1+18:1)}{16:0}$ |
| 1 | MG1655 | 28.64±0.25 | 22.14±0.23 | 30.05±0.29 | 6.50±0.05 | 3.30±0.04 | 1.7 |
| 2 | $\Delta fadR$ | 21.73±0.04 | 11.4±0.47 | 40.58±0.34 | 11.75±0.45 | 3.92±0.04 | 0.8 |
| 3 | $\Delta relA \Delta spoT$ | 28.37±0.27 | 17.4±0.26 | 33.0±0.21 | 7.4±0.27 | 3.71±0.03 | 1.4 |
| 4 | $\Delta relA \Delta spoT \Delta fadR$ # | 20.30±1.2 | 8.18±0.25 | 39.94±0.35 | 15.18±0.57 | 5.0±0.54 | 0.7 |
| 5 | $\Delta relA \Delta spoT \Delta fadR/pCA24N$ # | 18.8±0.70 | 8.84±0.34 | 43.5±1.21 | 12.7±0.56 | 4.39±0.11 | 0.6 |
| 6 | $\Delta relA \Delta spoT \Delta fadR/pCAgnsA$ | 29.3±0.23 | 23.8±1.9 | 22.31±0.95 | 5.54±0.29 | 5.35±0.33 | 2.4 |
| 7 | $\Delta relA \Delta spoT \Delta fadR/pCAfabB$ | 26.02±0.15 | 11.01±0.1 | 37.5±0.47 | 10.3±0.18 | 3.28±0.04 | 1 |
| 8 | $\Delta relA \Delta spoT \Delta fadR/pBAD33$ # | 18.70±0.16 | 7.14±0.63 | 42.12±1.42 | 14.65±0.14 | 5.63±0.26 | 0.6 |
| 9 | $\Delta relA \Delta spoT \Delta fadR/pBADfabA$ | 27.09±0.17 | 11.30±0.49 | 35.13±0.85 | 11.61±0.34 | 4.48±0.31 | 1.1 |

To sum up, expression of *gnsA*, *fabA* or *fabB* from plasmid rescued the growth defect of (p)ppGpp deficient Δ*fadR* strains (Fig. 2) and also to varying extents increased the relative proportion of unsaturated fatty acids in Δ*relA* Δ*spoT* Δ*fadR* strain (Table 1, last column). From this data and the finding that growth defect of (p)ppGpp deficient Δ*fadR* strains was rescued by an increase in growth temperature (Fig. 1) we made the following hypothesis. The growth defect of (p)ppGpp deficient Δ*fadR* strains follows from the decrease in membrane fluidity due to decrease in unsaturated fatty acid content and an increase in the growth temperature rescued growth by increasing membrane fluidity. The experiments described below were performed to test this hypothesis.

### Supplementation of palmitoleic acid (16:1) rescued the low temperature growth defect of (p)ppGpp deficient Δ*fadR* mutant

To test the hypothesis, we studied the effect of membrane associated fatty acids on growth of (p)ppGpp deficient Δ*fadR* strains. In *E. coli*, palmitoleic acid (16:1), cis-vaccenic acid (18:1) (both unsaturated fatty acids) and palmitic acid (16:0) (saturated fatty acid), are the primary constituents of membrane phospholipids (18). 16:1, but not 16:0, 18:1 or Brij-58 (solvent) rescued the plating defect of Δ*relA* Δ*fadR* strain at 25°C (Fig. 3A) and growth of Δ*relA* Δ*spoT* Δ*fadR* strain at 30°C (white colonies in Fig. 3B, panels i to iv). Growth rescue by 16:1 supplementation was in concordance with the growth rescue through expression of *gnsA*, *fabA* or *fabB,* all of which increase the 16:1 content (Fig. 2 and Table 1). That 18:1 did not rescue similar to 16:1 was unexpected. However, comparatively weaker rescue was observed with 18:1 in plating efficiency experiments with Δ*relA* Δ*fadR* strain, but this was not reproducible. 18:1 supported slow growth of Δ*relA* Δ*spoT* Δ*fadR* strain at 30°C (small white colonies in Fig. S2). The data showed, rescue by 16:1 was better than by 18:1.

**Figure 3.**
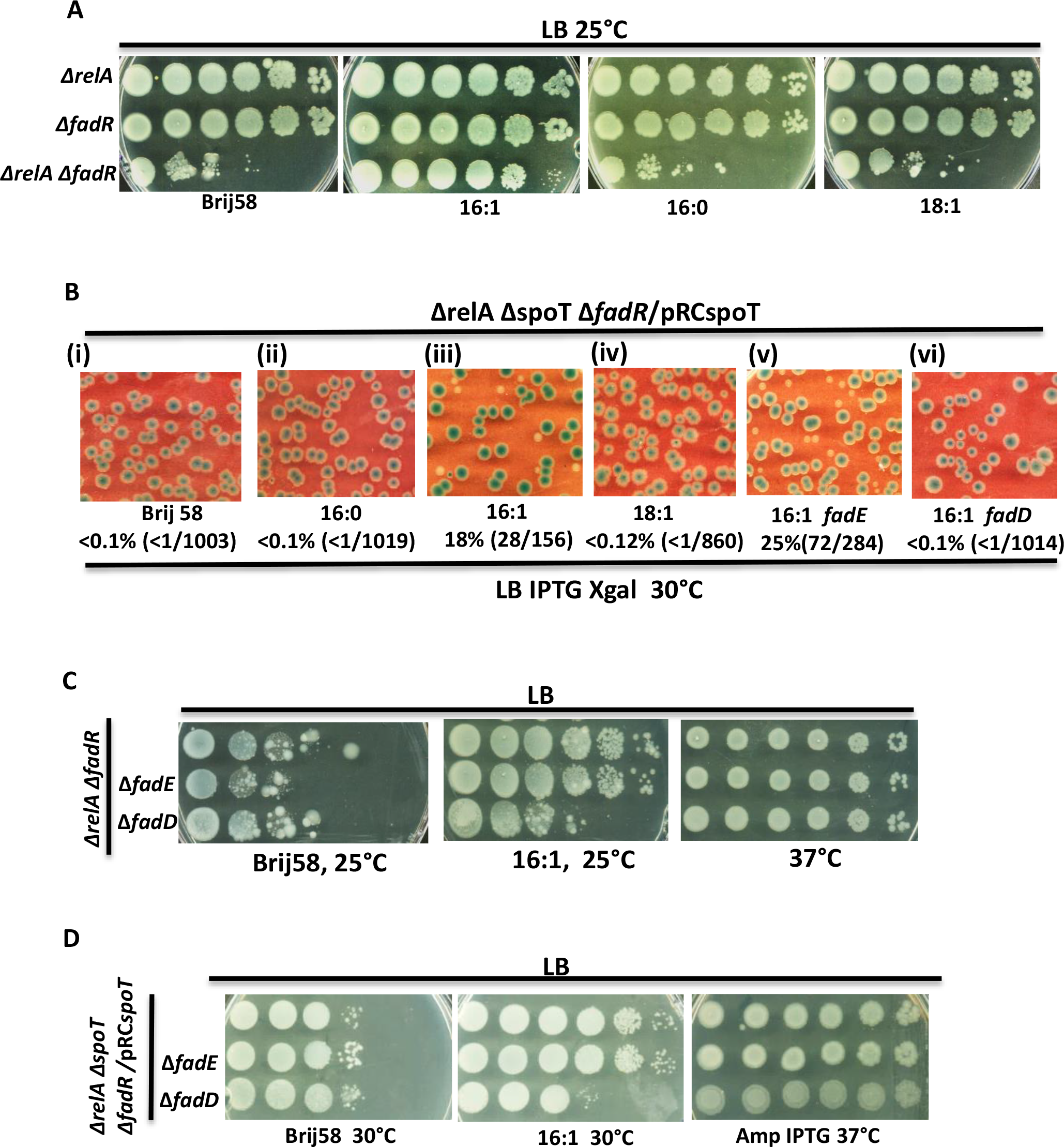
Membrane lipid synthesis with palmitoleic acid (16:1) rescued the low temperature growth defect of (p)ppGpp deficient Δ*fadR* mutant. Strains whose relevant genotypes are mentioned were cultured to stationary phase in LB medium or LB medium containing IPTG at 37⁰C, washed, serially diluted with minimal A medium and spotted on LB agar plates (A, C, D). Suitable dilutions were spread on LB agar plates containing IPTG and X-Gal to estimate the frequency of pRCspoT plasmid loss by quantification of blue/white colonies. The incubation temperature, fatty acids and antibiotics supplemented to the plates are mentioned besides the panels. The percentage of white colonies and the number of white colonies out of total colonies counted (blue+white) are shown below each panel. The strains are, Δ*relA* (VS3), Δ*fadR* (VS11), Δ*relA* Δ*fadR* (VS33), Δ*relA* Δ*spoT* Δ*fadR*/pRC*spoT* (VS7), Δ*relA* Δ*spoT* Δ*fadR* Δ*fadE* /pRC*spoT* (VS76), Δ*relA* Δ*spoT* Δ*fadR* Δ*fadD* /pRC*spoT* (VS75), Δ*relA* Δ*fadR* (VS159), Δ*relA* Δ*fadR* Δ*fadE* (VS210), Δ*relA* Δ*fadR* Δ*fadD* (VS211). All strains have the Δ*lacZYAI*::FRT allele

In *E. coli*, exogenous long chain fatty acids transported through the FadL/FadD system is converted to acyl-coA and catabolized through β-oxidation or the acyl chain is directly incorporated into the membrane phospholipid (19, 32). We examined the individual role of *fadD,* encoding long-chain-fatty-acid-CoA ligase required for transport and *fadE* encoding acyl-CoA dehydrogenase, required for degradation of exogenous long chain fatty acids in the growth rescue by 16:1. The rescue of Δ*relA* Δ*fadR* mutant by 16:1 at 25°C, was abolished in the presence of Δ*fadD* mutation but not Δ*fadE* mutation (Fig. 3C) and the same was seen for Δ*relA* Δ*spoT* Δ*fadR*/ pRC*spoT* strain at 30°C (Fig. 3D). Growth of Δ*relA* Δ*spoT* Δ*fadR* strain at 30°C in the presence of 16:1 (white colonies in Fig. 3B, panel iii) was abolished by Δ*fadD* mutation but not Δ*fadE* mutation (Fig. 3B, panels v & vi). These results indicated uptake and incorporation of 16:1 into membrane but not its degradation through β-oxidation was the likely reason for growth rescue.

### Δ*relA* Δ*spoT* strain was sensitive to reduction in proportion of 16:1+18:1 relative to 16:0

Inactivation of FadR, in addition to inhibition of unsaturated fatty acid biosynthesis influences overall fatty acid metabolism (17, 21). To understand role of (p)ppGpp in growth of strains specifically deficient in unsaturated fatty acid biosynthesis, we studied consequences of inhibiting enzymes required for unsaturated fatty acid biosynthesis in the Δ*relA* Δ*spoT* (*fadR*^+^) strain.

The essential *fabA* gene codes for 3-hydroxydecanoyl-ACP dehydrase, which catalyses the first committed step in the pathway of unsaturated fatty acid biosynthesis (19). Strains with the temperature sensitive *fabA2*(Ts) lesion exhibits increasing growth defect with increase in temperature due to progressive impairment of the enzyme’s catalytic activity (33). Under our experimental conditions, the *fabA2*(ts) allele conferred growth defect in an otherwise wild type strain at 42⁰C but not 37⁰C or 30⁰C. In the isogenic Δ*relA* Δ*spoT* background, we observed growth inhibition at 37⁰C and 42⁰C and not at 30⁰C (Fig. 4A). Assuming increase in growth temperature inhibited the catalytic activity of *fabA2* encoded protein equally in both strains, the result suggested, the (p)ppGpp deficient strain could be more sensitive than wild type to decrease in UFA concentration or decrease in proportion of UFA relative to SFA.

**Figure 4.**
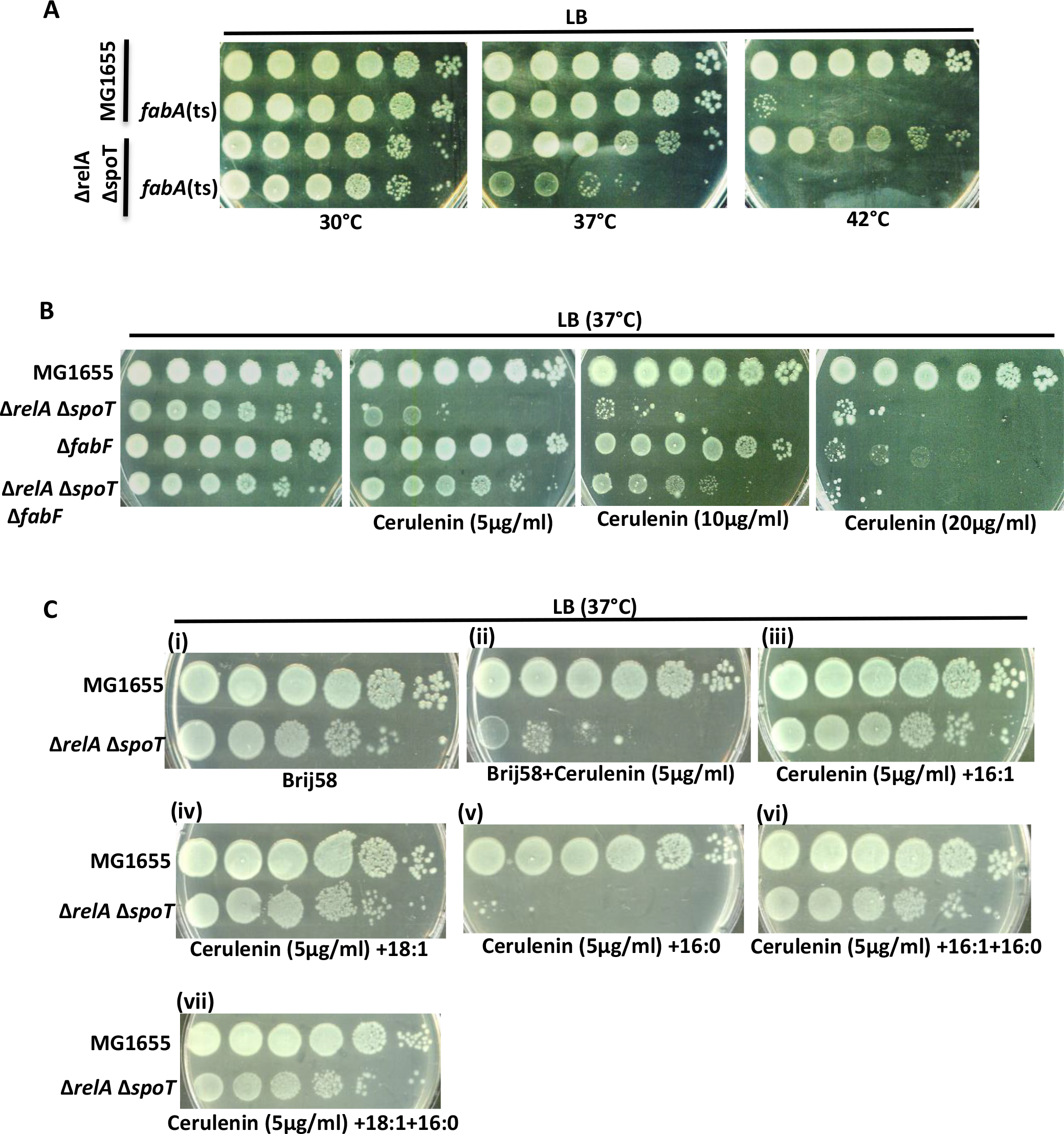
The (p)ppGpp deficient Δ*relA* Δ*spoT* strain was hypersensitive to perturbations that restrict unsaturated fatty acid biosynthesis. Strains whose relevant genotypes are mentioned were cultured to stationary phase in LB medium at 30⁰C (strains with *fabA2*(ts) and isogenic *fabA*^+^ alleles) or 37⁰C, washed, serially diluted with minimal A and spotted on LB agar plates. The incubation temperature, fatty acid supplemented and final concentration of cerulenin in the plates are mentioned besides the relevant panels. In case of Strains used are, MG1655 (VS2), MG1655 *fabA2*(ts) (VS155), Δ*relA* Δ*spoT* (AN120, white colony), Δ*relA* Δ*spoT fabA2*(ts) (VS157, white colony), Δ*fabF* (VS473), Δ*relA* Δ*spoT* Δ*fabF* (VS474, white colony). All strains have the Δ*lacZYAI*::FRT allele.

Type I and type II fatty acid synthase systems of prokaryotes and eukaryotes are inhibited by the antibiotic cerulenin, which covalently attaches to the active site cysteine of β-ketoacyl-ACP synthases and inactivates the enzyme (34). In *E. coli*, cerulenin inhibits FabB (3-ketoacyl-ACP synthase I) and FabF (3-ketoacyl-ACP synthase II), but FabB which is essential for synthesis of unsaturated fatty acid was more sensitive to the antibiotic (30, 35). Since FabF supports saturated fatty acid biosynthesis, and is more resistant to cerulenin than FabB, the continued synthesis of saturated fatty acid in presence of low concentration of cerulenin leads to relative decrease in the unsaturated fatty acids over saturated fatty acids in bulk lipid (35). Unsaturated fatty acid biosynthesis was more sensitive to cerulenin in *B. Subtilis* (36), suggesting this feature may be widely conserved in bacteria.

Cerulenin inhibited the growth of wild type strain when present at 80µg/ml (Fig. S3) and the Δ*fabF* mutant at 20µg/ml (Fig. 4B). The greater sensitivity of *ΔfabF* mutant can be due to the more cerulenin sensitive FabB enzyme solely carrying out long chain fatty acid synthesis. Interestingly, the Δ*relA* Δ*spoT* strain was hypersensitive to cerulenin – even more sensitive than *ΔfabF* mutant (Fig.4B) and this was rescued by supplementation of 16:1 or 18:1, but not 16:0 (Fig. 4C, panels i to v) – the latter appeared to enhance the cerulenin sensitivity (compare panels ii & v). The rescue by 16:1 or 18:1 was unaffected in the presence of 16:0 (Fig. 4C, vi and vii). These results are consistent with the idea that cerulenin hypersensitivity resulted from UFA deficiency.

It was possible, reduced expression of FabB/FabF (enzymes required for UFA synthesis) in the Δ*relA* Δ*spoT* strain was responsible for the cerulenin hypersensitivity. However, inactivation of FabF partially rescued the hypersensitivity phenotype (Fig. 4B), indicating FabF activity contributed to the hypersensitivity. It is unlikely, hypersensitivity was due to FabF mediated 18:1 synthesis (from 16:1), because (i) 18:1 rescued the phenotype and (ii) 16:1 synthesis would also be restricted as FabB (essential for 16:1) is more sensitive than FabF to cerulenin (30, 35). This leaves the possibility of loss of FabF mediated saturated fatty acid biosynthesis being responsible for partial rescue of cerulenin hypersensitivity, and suggested the strain lacking (p)ppGpp could be sensitive to an increase in proportion of SFA (primarily16:0) over UFA (16:1+18:1) in membrane and consequent decrease in fluidity. Consistent with this idea, growth improved at the higher temperature of 42⁰C (Fig. S4), which increases fluidity (37).

In *E. coli*, the relative proportion of cis-vaccenic acid (18:1) to palmitic acid (16:0), increases with decrease in growth temperature (thermal regulation) (38). The lower transition temperature of unsaturated fatty acids keeps the membrane fluid (37). Work from Cronan lab showed, thermal regulation of fatty acid composition arises from the intrinsic temperature sensitivity of FabF enzyme, and therefore more efficient synthesis of cis-vaccenic acid (18:1) at low temperature (39 & references therein). Unlike in Δ*fadR* background, (p)ppGpp deficiency in Δ*fabF* background did not confer growth defect at 25⁰C or 30⁰C (Fig. S5, compare with Figs. 1C and 1D). Importantly, while the *fadR* mutation lowers the proportion of UFA over SFA ∼2-fold in the membrane (Table 1, 31), *fabF* mutation does not alter this proportion, because inhibition of 18:1 synthesis increases 16:1 content (40, 41). Taken together the results supports the idea, decrease in proportion of UFA over SFA caused the low temperature specific growth defect of (p)ppGpp deficient Δ*fadR* strains.

To find out if (p)ppGpp was required for adapting to growth conditions that change membrane fatty acid composition, we tested the effect of long and medium chain fatty acids on the growth of wild type and Δ*relA* Δ*spoT* strains. Palmitic acid (16:0) severely inhibited growth of Δ*relA* Δ*spoT* mutant but not isogenic wild type strain (Fig. 5A, panels i and ii). Lauric acid (12:0), myristic acid (14:0), palmitoleic acid (16:1) or cis-vaccenic acid (18:1) did not affect growth of both strains (panel iii to vi). Supplementation of 16:1 or 18:1 rescued the growth inhibition conferred by 16:0 (Fig. 5A, panel vii and viii). Since 16:0, 16:1 and 18:1 are the fatty acids that make up the membrane lipids, the result implicates the increase in relative proportion of 16:0 (over 16:1 and 18:1) being responsible for the growth inhibition of Δ*relA* Δ*spoT* strain. Using the *ΔfadE* allele, we confirmed degradation of 16:0 was not required for its growth inhibitory effect and degradation of 16:1 or 18:1 was not required for their growth rescue in the (p)ppGpp deficient strain. As expected, FadD mediated uptake of 16:0 was required for its growth inhibitory effect (Fig. 5B).

**Figure 5.**
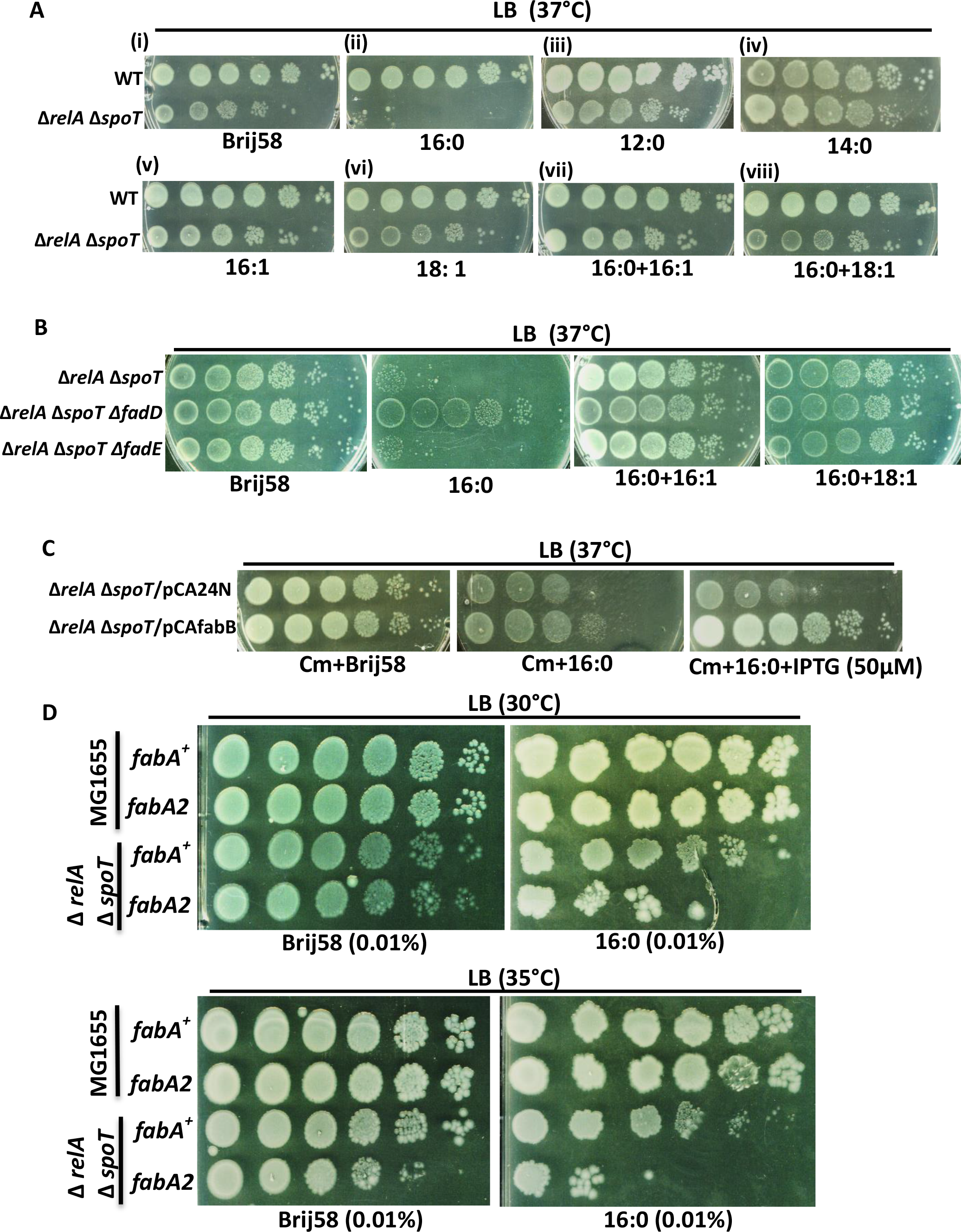
Relative proportion of 16:0 versus (16:1+18:1) in membrane was the critical determinant of growth in Δ*relA* Δ*spoT* strain. Strains whose relevant genotypes are mentioned were cultured to stationary phase in LB medium at 30⁰C (strains with *fabA2*(ts) and isogenic *fabA*^+^ alleles) or 37⁰C or without antibiotics, washed, serially diluted with minimal A medium and spotted on LB agar plates (A, B and C). The incubation temperature, the fatty acid supplemented, and antibiotics used for plasmid selection are mentioned beside the panels. MG1655 (VS2), Δ*relA* Δ*spoT* (AN120, white colony), Δ*relA* Δ*spoT*/pCA24N (VS479), Δ*relA* Δ*spoT*/pCAfabB (VS478), Δ*relA* Δ*spoT* Δ*fadD* (VS418, white colony), Δ*relA* Δ*spoT* Δ*fadE* (VS419, white colony), MG1655 *fabA2*(ts) (VS155), Δ*relA* Δ*spoT fabA2*(ts) (VS157, white colony).

Acyl-CoA intermediates are produced concomitant with uptake of exogenous fatty acids, while endogenously synthesized fatty acids make acyl-ACP intermediates and both intermediates can be used for synthesis of membrane lipids (19, 42). Over-expression of *fabB* rescued the growth inhibition conferred by 16:0 (Fig. 5C). This indicated, increasing the intracellular concentration of 16:1-ACP and 18:1-ACP can counteract the growth inhibition caused by 16:0-CoA by competing for incorporation into membrane lipids. Finally, to confirm that a relative increase in content of 16:0 over UFAs was the critical determinant for growth inhibition by 16:0, we reduced the concentration of 16:0 in growth medium till it was no longer growth inhibitory (from 0.05% to 0.01%) and reinstated growth inhibition by lowering unsaturated fatty acid synthesis using the *fabA2*(ts) allele (Fig. 5D). It is known, the *fabA2*(ts) encoded protein has less catalytic activity than wild type FabA at the growth temperatures used in the experiment (35⁰C or 30⁰C) (33). Taken together, the results provide strong evidence that (p)ppGpp was required for growth when the proportion of unsaturated fatty acids (16:1+18:1) was reduced relative to saturated fatty acid (16:0) in the membrane.

### Growth defect of strains deficient in (p)ppGpp and unsaturated fatty acids was associated with cell division defect and lysis and rescued by *ftsQAZ* expression

When cultured at 25°C in broth from permissive growth condition, the Δ*relA* Δ*fadR* strain but not isogenic Δ*relA* or Δ*fadR* strains exhibited growth arrest (Fig. S6A). Similarly, the Δ*relA* Δ*spoT* Δ*fadR*/pRC*spoT* strain exhibited growth defect when cultured at 30°C without IPTG, but not in the presence of IPTG (Fig. S6B). FACS analysis following propidium iodide staining showed the growth arrested cells had significantly higher membrane integrity defects as compared to isogenic strains (Fig. S7). Since (p)ppGpp is implicated in regulation of cell division and membrane biogenesis (10, 11, 12, 13), we examined the morphology of cells undergoing growth arrest and compared it to cells of isogenic strains that did not exhibit growth defect. Live cell imaging was performed to follow cell growth and division in the (p)ppGpp deficient Δ*fadR* strains following shift from permissive to non-permissive growth conditions. Isogenic strains cultured under identical growth conditions served as controls. Starting from 2 to 4 cells, images were captured at periodic intervals as the cells underwent several rounds of division (movies 1 to 9).

In LB medium at 25°C, growth and cell division was largely similar between wild type (VS2) (movie 1) and Δ*fadR* (VS11) (Movie 2) cells. The cells of Δ*relA* (VS3) (Movie 3) strain grew longer before undergoing cell division. In case of Δ*relA* Δ*fadR* cells (VS159), after few rounds of division, cells started to filament and several of them underwent lysis (Movie 4) which is consistent with the growth inhibition observed in the strain. In LB medium at 30°C, growth and cell division were largely similar between the wild type (VS2) (movie 5) and Δ*fadR* (VS11) (movie 6) cells. In Δ*relA* Δ*spoT* strain (AN120, white colony), cells grew longer (as compared to wild type or Δ*fadR* cells) before cell division and few of them formed filaments (movie 7). There was more heterogeneity in cell size. This is consistent with data reported in other studies (12, 43, 44). The Δ*relA* Δ*spoT* Δ*fadR*/pRC*spoT* strain cultured without IPTG, that is under (p)ppGpp depleted condition formed distinctly long/filamented cells that underwent lysis (VS6) (movies 8 & 9).

Since cell of the (p)ppGpp deficient Δ*fadR* strains showed cell division defect before lysis, we studied the effect of over-expressing cell division proteins in these strains. The formation of FtsZ ring is at the heart of bacterial cytokinesis and provides the platform for assembly of divisome complex needed for cell septation (45). Since *ftsQAZ* genes form an operon, and expression of *ftsZ* is driven from promoters located within *ftsQ* and *ftsA*, a plasmid construct having all three genes (pCL*ftsQAZ*) was used to study its effect on the growth of (p)ppGpp deficient Δ*fadR* strains.

At 30°C, pCL*ftsQAZ* but not the vector rescued the plating defect of Δ*relA* Δ*spoT* Δ*fadR*/ pRC*spoT* strain after IPTG withdrawal (Fig. 6A (i)) and supported growth of Δ*relA* Δ*spoT* Δ*fadR* strain at 30°C (white colonies in Fig. 6A (ii)). Live cell imaging showed *ftsQAZ* expression from plasmid rescued the cell filamentation and lysis phenotypes seen under the (p)ppGpp depleted growth condition in this strain (VS452) (movie 10), while the phenotypes persisted in the presence of vector (VS451) (movie 11). *ftsQAZ* expression rescued the hypersensitivity of Δ*relA* Δ*spoT* strain to cerulenin (Fig. 6B) and its sensitivity to palmitic acid (16:0) in growth medium (Fig. 6C). These results strongly suggest the growth inhibition observed under these conditions (decrease in UFA content) was due to impaired cell division.

**Figure 6.**
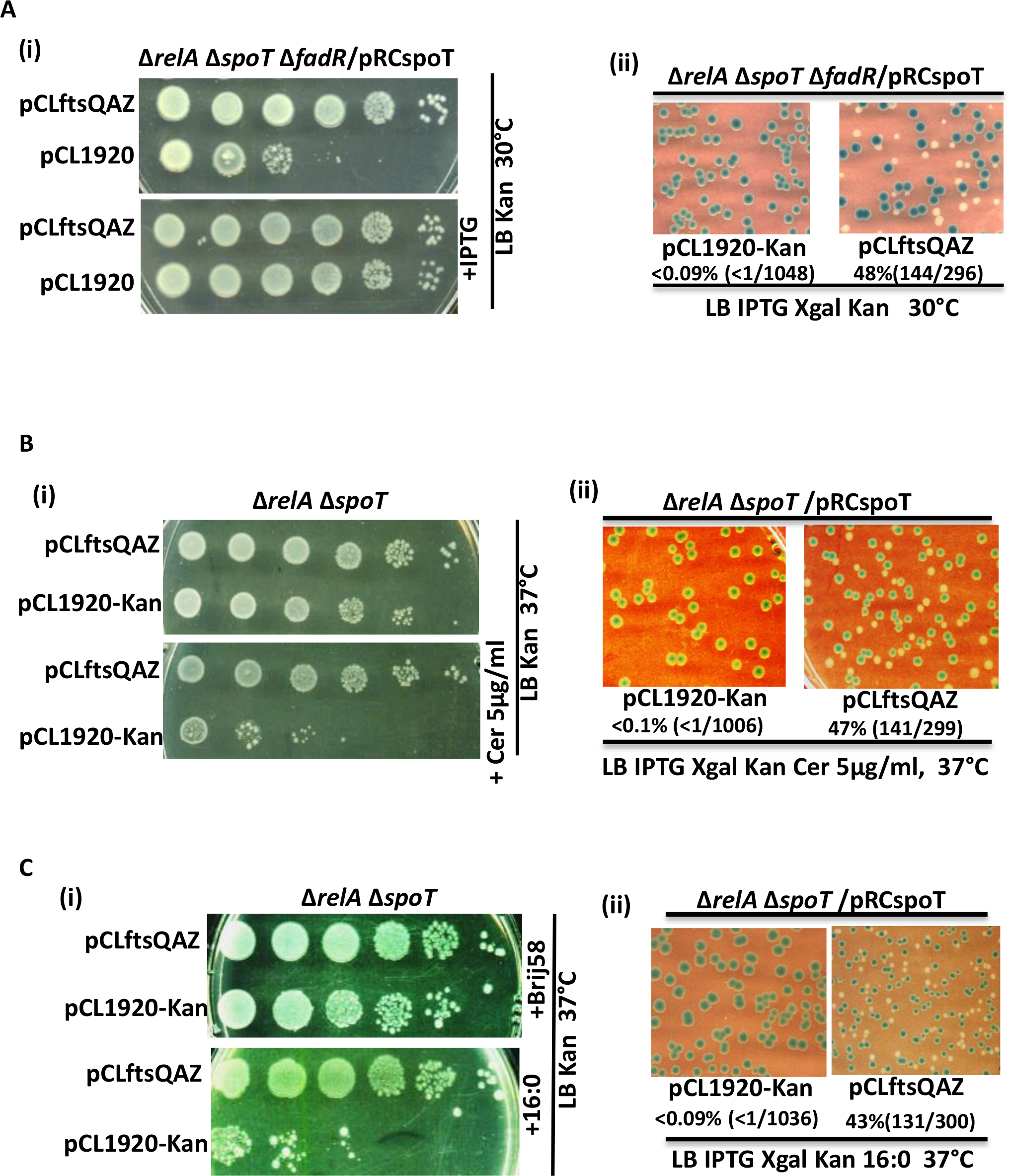
Expression of *ftsQAZ* rescued the growth defect of (p)ppGpp deficient strains with reduced unsaturated fatty acid synthesis. Strains whose relevant genotypes are mentioned were cultured to stationary phase in LB medium at 37⁰C with antibiotics and IPTG as appropriate, washed, serially diluted in minimal A medium and spotted on LB agar plates [A(i), A(ii), B(i) and C(i)], or suitable dilutions spread on LB agar plates containing IPTG and X-Gal to estimate the frequency of pRCspoT plasmid loss by quantifying the number of blue/white colonies [A(ii), B(ii), C(ii)]. The incubation temperature, fatty acids and antibiotics supplemented to the plates are mentioned besides the panels. The percentage of white colonies obtained, number of white colonies out of total colonies used for the calculation are also shown below appropriate panels. Strains used are, Δ*relA* Δ*spoT* Δ*fadR*/ pRC*spoT* /pCL1920 (VS451), Δ*relA* Δ*spoT* Δ*fadR*/ pRC*spoT* /pCLQAZ (VS452), Δ*relA* Δ*spoT*/ pRC*spoT* /pCL1920 (VS469), Δ*relA* Δ*spoT*/ pRC*spoT* /pCLQAZ (VS470), Δ*relA* Δ*spoT*/pCL1920 (VS469, white colony), Δ*relA* Δ*spoT*/pCLQAZ (VS470, white colony).

## Discussion

The stringent response provoked by the guanine nucleotide analogues (p)ppGpp alter the expression and activity of large number of genes and proteins and exert global effect on cell physiology (3, 6, 46). Classical view of stringent response primarily based on studied in *E. coli* has been that, following starvation (p)ppGpp accumulates to cause extensive re-programming by altering gene-expressions leading to growth inhibition and preservation of cell viability. Another view has been, depending of the concentration of (p)ppGpp different cellular processes are regulated in a continuum. The second view has gained prevalence as studies in *E. coli* and other bacteria have revealed a large number of targets with different binding affinity for (p)ppGpp (47 and references therein). By analyzing binding affinities, inhibitory constants and EC50/IC50 values for targets of (p)ppGpp a scheme of prioritization in (p)ppGpp mediated direct regulation has been proposed (47). In the backdrop of the second view, we have looked for (p)ppGpp driven regulations necessary for growth in response to specific genetic or environmental perturbation. This could identify regulations (direct or indirect) necessary for adaptation to specific stress (in contrast to regulations whose biological significance in the context of the stress may not be immediately obvious).

One way to identify such (p)ppGpp dependent adaptations could be through genetic screens to uncover mutations that confer synthetic lethal (or synthetic growth defect) phenotypes in the (p)ppGpp deficient background (23, 48, 49, 50). In this study, we characterized the synthetic growth defect arising from (p)ppGpp deficiency together with partial starvation for unsaturated fatty acids. The study has revealed the existence of (p)ppGpp dependent adaptive response essential for functioning of cell division machinery under conditions of reduced membrane fluidity, which to our knowledge has not been recognized previously.

Arguably, homeoviscous adaptation in response to temperature change is the best studied membrane adaptive process and primarily involves maintenance of fluidity by altering lipid composition. The cellular consequences that follow from loss of homeoviscous adaptation in *E. coli* and *B. subtilis* was addressed in a recent study (51). This study, in agreement with previous study by Cronan & Gelmann (33) found 10-15% UFAs was essential to support growth in *E. coli*. At this point, large-scale lipid phase separation associated with loss of several essential membrane functions (including cell division in case of *E. coli*) was observed (51). Our results show in the (p)ppGpp deficient strain, the process of cell division was inhibited much before UFAs reached these low levels. Under our experimental condition, when the UFAs, which are 56% of total in the wild type strain drops to 37% of total in the Δ*fadR* strain (Table 1), growth is almost normal (movies 5 & 6). However, a further decrease to 32% in the (p)ppGpp depleted Δ*fadR* mutant led to the disruption of cell division and lysis (movies 8 & 9). This clearly indicates a (p)ppGpp dependent adaptive process was required for cell division before UFAs are reduced to 10-15% resulting in growth arrest from large scale lipid phase separation but without membrane rupture.

Since (p)ppGpp accumulates following fatty acid starvation (14), identification of (p)ppGpp dependent regulation to survive loss of homeoviscous adaption was not entirely unexpected. However, this has to considered in the context of current understanding of stringent response to fatty acid starvation. RelA (52) and SpoT (14, 53) mediated synthesis of (p)ppGpp was reported following severe fatty acid starvation that produces growth inhibition (classical stringent response). Under our experimental conditions (Δ*fadR* mutant cultured in LB medium) the mild starvation, primarily for UFAs could signal (p)ppGpp synthesis and set a higher steady state basal level. Our results show, when homeostatic adaptation was impaired in the Δ*fadR* mutant, both RelA and SpoT mediated (p)ppGpp synthesis contributes to the increase in basal level and this was necessary for cell division depending on the extent of membrane fluidity loss. Both RelA and SpoT dependent (p)ppGpp synthesis was necessary at 25⁰C (Fig. 1A), SpoT dependent synthesis was sufficient at 30⁰C and 37⁰C (Figs. 1A & 1C), and (p)ppGpp was not required at 42⁰C (Fig. 1D).

Since cell division in all life forms is an intrinsically membrane associated process, it being sensitive to reduction in membrane fluidity may be expected. In *E. coli*, the assembly of divisome initiates with the formation of FtsZ filaments that attach to the cytoplasmic membrane by the binding of conserved C-terminal peptide portion of FtsZ with the membrane anchor proteins FtsA and ZipA (45). It is possible, membrane fluidity plays a critical role in the interaction of anchor proteins with cytoplasmic membrane or in later step(s) in the divisome assembly, especially under (p)ppGpp deficient conditions. Further studies are needed to understand details of the mechanism by which (p)ppGpp supports divisome function under conditions of reduced fluidity.

## Materials & Methods

### Media and chemicals

Media used in this study are, LB (0.5% yeast extract, 1% tryptone, and 1% NaCl), LB – NaCl (0.5% yeast extract, 1% tryptone), minimal A (75), and PBS (phosphate buffered saline). For selection, antibiotics were used at final concentration of 25μg/ml kanamycin (Kan), 15μg/ml chloramphenicol (Cm), 50 μg/ml ampicillin (Amp), 10 μg/ml tetracycline (Tet), and 25 μg/ml spectinomycin (Spec). 5-Bromo-4-chloro-3-indolyl-D-galactoside (X-gal) was used at final concentration of 50μg/ml, isopropyl-D-thiogalactopyranoside (IPTG) at 1mM unless stated otherwise and propidium iodide at 1µg/ml. For preparation of fatty acids solutions, a 10% solution of Brij58 was made in water and used to prepare fatty acid stocks at a concentration of 2% and neutralised with KOH. Boiling water bath was used to dissolve fatty acids. All the fatty acids were used at final concentration of 0.05% unless stated otherwise. All fatty acids used in the study were purchased from Sigma Aldrich.

### Bacterial Strains, Plasmids and Genetic procedures

All strains were constructed in the MG1655 background. Phage P1 transduction, cloning and other techniques were carried out using standard procedures (54, 55). Strains, plasmids and primers used in the study are given in Table S1 of the supplemental material. Gene deletions were obtained from the Keio collection (56), and pCP20 plasmid that codes for FLP recombinase was used to remove the Kan cassette whenever necessary (57). Appropriate plasmid clones sourced from ASKA collection were verified and used for over-expression of *E. coli* genes (58). The F derived single-copy plasmid pRC7 (24) was used to create the plasmids pRC*spoT* and pRCspec-*spoT* (23). In these plasmids the *spoT* gene with native RBS and start codon is expressed from the *lac* promoter and placed immediately upstream of *lacZ* gene. The *lacZ* gene is also expressed from the *lac* promoter. *fabA* gene with its own Shine-Dalgarno sequence was PCR amplified from MG1655 genome using the primer pairs JGABkpnfabAFP and JGABfabAH3RP and cloned into the *Kpn*I and *Hin*dIII sites of modified pBAD33 vector (pBAD33m) to generate pBAD*fabA*. Spectinomycin resistance cassette (*aadA* gene) of pCL1920 and the pMN8 plasmid (59) carrying the *ftsQAZ* genes was replaced with kanamycin resistance cassette (*npt* gene) through recombineering (60) using primers JGABKanpKD13FP and JGABKanpKD13RP. The Δ*relA* Δ*spoT*/pRC*spoT* strain maintained in LB medium containing ampicillin and IPTG was used to obtain the Δ*relA* Δ*spoT* strain by streaking on LB IPTG X-plates and scoring for white colonies. Viability of cells was scored by spotting 10µl of serial 10-fold dilutions from overnight cultures that were washed and diluted with minimal A medium on appropriate plates and incubated for 56-72 hours at 25°C, 36-48 hours at 30°C and 16-24 hours at 37°C. The growth rate and growth defect of strains was monitored in broth by measuring absorbance at A600.

### Plasmid segregation assay

The “blue-white” plasmid segregation assay can be used to investigate synthetic lethality/growth defect between mutations. The assay is based on the rationale that the essentiality of plasmid encoded gene function would stabilize an otherwise unstable plasmid. The single-copy pRCspoT plasmid, which is unstable in the absence of selection was used to investigate the essentiality of *spoT* gene function in different genetic backgrounds. All strains used in this assay carried the Δ*lacZYAI*::FRT allele on the chromosome and therefore presence of pRCspoT plasmid can be visually scored using the plasmid encoded “*lac+*” phenotype in plates containing IPTG and X-gal. Colonies with cells retaining pRCspoT are blue and while those with cells that have lose it are white. Colony can be sectored if plasmid loss happens during the growth of colony. SpoT function is normally not essential for growth in the Δ*relA* Δ*spoT* background, therefore, stabilisation of pRCspoT, that is, occurrence of only blue colonies in the Δ*relA* Δ*spoT* genetic background indicated (p)ppGpp synthesis was indispensable for growth of the strain. Strains used for this assay were cultured overnight without selection for pRCspoT plasmid at 37°C, washed once and serially diluted with minimal A medium and spread on LB IPTG X-Gal plates to obtain 100-200 colonies per plate. Plates were incubated for 24 – 36 hours at 37°C or 42°C, 48 – 72 hrs at 30°C, and the proportion of white colonies calculated as the number of white colonies over total (blue colonies + sectored colonies + white colonies).

### Live cell imaging

All strains were cultured overnight in LB medium at 37°C. Overnight cultures of MG1655, Δ*relA*, Δ*fadR* and Δ*relA ΔfadR* strains were diluted 200-fold into LB medium and cultured at 25°C. MG1655, Δ*fadR* and Δ*relA* Δ*spoT* strains were diluted 500-fold into LB medium and cultured at 30°C. The Δ*relA* Δ*spoT ΔfadR*/pRC*spoT* strain cultured overnight in the presence of ampicillin and IPTG was washed twice with LB and diluted 500- fold into LB medium. When pCLKan1920 or pCLKan*ftsQAZ* was present in the Δ*relA* Δ*spoT ΔfadR*/pRC*spoT* strain, kanamycin was added to LB medium at all stages. All cultures were grown to an A600 of ∼ 0.1 and 10 μl was spread in the centre of cell culture dish (CELL VIEW) and allowed to dry. Before complete dehydration, 400 μl of warm molten LB agar was spread over the culture film and allowed to dry for 5 minutes. The culture dish was closed, sealed with parafilm tape, incubated at appropriate temperature for 30 minutes to 1 hour and then placed within the chamber of LEICA TCS SP8 DMi8 microscope preset to the required growth temperature and imaged for 4 to 6 hours with pictures captured every 2 minutes.

### Determination of fatty acid profile

For each strain, three independent overnight broth cultures (growth at 37⁰C) were made following inoculation from three independent colonies. Cells were diluted 10^4^-fold into fresh broth, cultured at 30°C to ∼ 0.2 A600 and harvested from 100 ml. The cultures were placed on ice to bring the temperature between 0°C to 4°C, washed twice with pre-chilled LB and cells pelleted at 4°C were frozen using liquid nitrogen. For selection of plasmids (other than pRC*spoT*) and maintenance of gene expression, appropriate antibiotics and inducers were included at all stages of growth. In case of pRC*spoT*, ampicillin and IPTG was included to overnight culture but not after sub-culture. When cells were harvested under these growth conditions, strains that exhibit growth defect on LB agar plates at 30°C, such as, Δ*relA* Δ*spoT ΔfadR*/pRC*spoT* (Fig. 1C), Δ*relA* Δ*spoT ΔfadR*/pRC*spoT*/ pCA24N (Fig. 2B), and Δ*relA* Δ*spoT ΔfadR*/pRC*spoT*/pBAD33m (Fig.2B) showed severe growth defect (data not shown). Fatty acid methyl ester (FAME) extraction and analysis was performed as previously described (61). Briefly, lipids were saponified in sodium hydroxide and methanol, methylated in acidified methyl alcohol, extracted in hexane and methyl tertiary butyl ether, and analyzed by using a gas chromatograph equipped with a flame ionization detector. The extraction efficiency of the protocol and authenticity of fatty acid peaks were verified using *Stenotrophomonas maltophilia* ATCC 13637T (with a known fatty acid profile) as a positive control. Peaks were identified based on retention time of a standard run under similar set of conditions, using the software and database (RTSBA6) of MIS (MIDI Inc., Newark, DE).

### Quantification of dead cells using flow cytometry

For each strain, three independent overnight broth cultures (growth at 37⁰C) were made following inoculation from three independent colonies. Whenever IPTG was present in the growth medium, cultures were washed twice with LB medium to remove IPTG and diluted 10^4^-fold into fresh LB medium at 30°C or directly diluted 10^3^-fold into LB at 25°C. 1 ml of early log phase cells were mixed with propidium iodide (1µg/ml) and incubated in the dark for 20 minutes. As expected, early log phase cultures of Δ*relA ΔfadR* and Δ*relA* Δ*spoT ΔfadR*/pRC*spoT* strains showed severe growth defect. All cultures were washed twice with PBS and used for flow cytometry in the BD-FACS Aria III platform. The BD-FACS Diva software (version 6.0) was used to analyse data from 20,000 cells in each preparation.

## ACKNOWLEDGMENTS

We thank Dr. Neetha Joseph (National Centre for Microbial Resources), Pune, for performing the FAME analysis. We acknowledge NBRP, Japan for strains and plasmids from keio collection and ASKA library and Coli Genetic Stock Center (CGSC) for the *fabA2*(Ts) strain. This work was funded by research grant to Centre of Excellence in Microbial Biology (Phase2) and BT/PR35733/BRB/10/1839/2019 to R.H from Department of Biotechnology (DBT), Government of India. V.S. is a recipient of CSIR-SRF fellowship.

## Conflict of Interest

We declare to have no conflict of interest.

## Author Contribution

VS and RH designed experiments, analysed data; VS performed experiments, wrote initial draft; RH wrote the manuscript

---

**Figure S1 - Complementation of *fabA2*(ts) mutant by pBAD*fabA*.** Strains whose relevant genotypes are mentioned were cultured to stationary phase in LB medium containing chloramphenicol at 30⁰C, washed, serially diluted with minimal A medium and spotted on LB agar plates with chloramphenicol. The incubation temperature and inducer (arabinose) added are indicated besides the panels. Strains used are *fabA2*(ts)/pBAD33m (VS484), *fabA2*(ts)/pBAD*fabA* (VS485).

**Figure S2 - Weak growth rescue of (p)ppGpp deficient** Δ***fadR* by cis-vaccenic acid (18:1).** The Δ*relA* Δ*spoT* Δ*fadR*/pRC*spoT* (VS6) strain was cultured to stationary phase in LB medium with 1mM IPTG, washed, serially diluted with minimal A medium and suitable dilution spread on LB agar plates containing IPTG and X-Gal and scored for blue and white colonies after 48 hrs of incubation at 30⁰C. Arrows point to tiny white colonies.

**Figure S3 - Differential sensitivity of wild type and** Δ***fabF* mutant to cerulenin.** The wild type (VS2) and Δ*fabF*::Kan (VS473) strains were cultured overnight to stationary phase in LB medium at 37⁰C, washed, serially diluted with minimal A medium and spotted on LB agar plates containing the indicated concentration of cerulenin and incubated at 37⁰C.

**Figure S4 – Increase in growth temperature alleviates cerulenin hypersensitivity of Δ*relA* Δ*spoT* strain.** The wild type (VS2) and Δ*relA* Δ*spoT* (AN120, white colony) strains were cultured to stationary phase in LB medium at 37⁰C, washed, serially diluted with minimal A medium and spotted on LB agar plates with or without 5µg/ml of cerulenin and incubated at the temperatures mentioned.

**Figure S5 – 18:1 synthesis is not required for growth of Δ*relA* Δ*spoT* strain at low temperature.** Strains whose relevant genotypes are mentioned were cultured to stationary phase in LB medium at 37⁰C, washed, serially diluted in minimal A medium and spotted. The growth medium and incubation temperature are indicated besides the panels. The strains are, MG1655 (VS2), Δ*fabF*::Kan (VS473), Δ*relA* Δ*spoT* (AN120, white colony) and Δ*relA* Δ*spoT* Δ*fabF* (VS474, white colony).

**Figure S6 - Growth defect of (p)ppGpp deficient Δ*fadR* strains in broth.** Strains whose relevant genotypes are indicated were cultured in LB medium to stationary phase at 37⁰C, IPTG and ampicillin was included for the Δ*relA* Δ*spoT* Δ*fadR*/pRC*spoT* strain. The Δ*relA* (VS3), Δ*fadR* (VS11), Δ*relA* Δ*fadR* (VS33) strains were diluted 10^3^-fold into LB medium and grown with shaking at 25⁰C. The Δ*relA* Δ*spoT* Δ*fadR*/pRC*spoT* (VS6) culture was washed and diluted 10^4^-fold into LB medium and grown with shaking at 30⁰C with or without IPTG. The optical density (A600) of the cultures were monitored and plotted against incubation time.

**Figure S7 - (p)ppGpp deficient Δ*fadR* strains exhibit membrane integrity loss during growth in LB medium.** Strains whose relevant genotypes are indicated were cultured to stationary phase in LB medium under permissive growth conditions and diluted as described in the legend to figure S6. The Δ*relA* Δ*spoT* (AN120, white colony) was diluted 10^4^-fold into LB medium. Cells were treated with propidium iodide and FACS analysis was carried out as mentioned under methods.

## Notes

### Competing Interest Statement

The authors have declared no competing interest.

## References

1. Bremer, H. and P. P. Dennis (2008). Modulation of Chemical Composition and Other Parameters of the Cell at Different Exponential Growth Rates. EcoSal Plus 3(1).

2. Atkinson, G. C., et al. (2011). The RelA/SpoT homolog (RSH) superfamily: distribution and functional evolution of ppGpp synthetases and hydrolases across the tree of life. PLoS One 6(8): e23479.

3. Potrykus, K. and M. Cashel (2008). (p)ppGpp: still magical? Annu Rev Microbiol 62: 35–51.

4. Sarubbi, E., et al. (1988). Basal ppGpp level adjustment shown by new spoT mutants affect steady state growth rates and *rrnA* ribosomal promoter regulation in Escherichia coli. Mol Gen Genet 213(2-3): 214–222.

5. Potrykus, K., et al. (2011). ppGpp is the major source of growth rate control in E. coli. Environ Microbiol 13(3): 563–575.

6. Liu, K., et al. (2015). Diversity in (p)ppGpp metabolism and effectors. Curr Opin Microbiol 24: 72–79.

7. Campos, M., et al. (2014). A constant size extension drives bacterial cell size homeostasis. Cell 159(6): 1433–1446.

8. Taheri-Araghi, S., et al. (2015). Cell-size control and homeostasis in bacteria. Curr Biol 25(3): 385–391.

9. Schreiber, G., et al. (1995). ppGpp-mediated regulation of DNA replication and cell division in Escherichia coli. Curr Microbiol 30(1): 27–32.

10. Joseleau-Petit, D., et al. (1999). Metabolic alarms and cell division in Escherichia coli. J Bacteriol 181(1): 9–14.

11. Buke, F., et al. (2022). ppGpp is a bacterial cell size regulator. Curr Biol 32(4): 870–877 e875.

12. Vadia, S., et al. (2017). Fatty Acid Availability Sets Cell Envelope Capacity and Dictates Microbial Cell Size. Curr Biol 27(12): 1757–1767 e1755.

13. Yao, Z., et al. (2012). Regulation of cell size in response to nutrient availability by fatty acid biosynthesis in Escherichia coli. Proc Natl Acad Sci U S A 109(38): E2561–2568.

14. Seyfzadeh, M., et al. (1993). spoT-dependent accumulation of guanosine tetraphosphate in response to fatty acid starvation in Escherichia coli. Proc Natl Acad Sci U S A 90(23): 11004–11008.

15. Battesti, A. and E. Bouveret (2006). Acyl carrier protein/SpoT interaction, the switch linking SpoT-dependent stress response to fatty acid metabolism. Mol Microbiol 62(4): 1048–1063.

16. Heath, R. J., et al. (1994). Guanosine tetraphosphate inhibition of fatty acid and phospholipid synthesis in Escherichia coli is relieved by overexpression of glycerol-3- phosphate acyltransferase (plsB). J Biol Chem 269(42): 26584–26590.

17. My, L., et al. (2013). Transcription of the Escherichia coli fatty acid synthesis operon fabHDG is directly activated by FadR and inhibited by ppGpp. J Bacteriol 195(16): 3784–3795.

18. Pulschen, A. A., et al. (2017). The stringent response plays a key role in Bacillus subtilis survival of fatty acid starvation. Mol Microbiol 103(4): 698–712.

19. Cronan, J. E., Jr. and C. O. Rock (2008). Biosynthesis of Membrane Lipids. EcoSal Plus 3(1).

20. Henry, M. F. and J. E. Cronan, Jr. (1992). A new mechanism of transcriptional regulation: release of an activator triggered by small molecule binding. Cell 70(4): 671–679.

21. My, L., et al. (2015). Reassessment of the Genetic Regulation of Fatty Acid Synthesis in Escherichia coli: Global Positive Control by the Dual Functional Regulator FadR. J Bacteriol 197(11): 1862–1872.

22. Zhang, F., et al. (2012). Enhancing fatty acid production by the expression of the regulatory transcription factor FadR. Metab Eng 14(6): 653–660.

23. Nazir, A. and R. Harinarayanan (2015). Inactivation of Cell Division Protein FtsZ by SulA Makes Lon Indispensable for the Viability of a ppGpp0 Strain of Escherichia coli. J Bacteriol 198(4): 688–700.

24. 24. Bernhardt, T. G. and P. A. de Boer (2004). Screening for synthetic lethal mutants in Escherichia coli and identification of EnvC (YibP) as a periplasmic septal ring factor with murein hydrolase activity. Mol Microbiol 52(5): 1255–1269.

25. Xiao, H., et al. (1991). Residual guanosine 3’,5’-bispyrophosphate synthetic activity of relA null mutants can be eliminated by spoT null mutations. J Biol Chem 266(9): 5980–5990.

26. Rock, C. O., et al. (1996). Increased unsaturated fatty acid production associated with a suppressor of the fabA6(Ts) mutation in Escherichia coli. J Bacteriol 178(18): 5382–5387.

27. Sugai, R., et al. (2001). Overexpression of yccL (gnsA) and ydfY (gnsB) increases levels of unsaturated fatty acids and suppresses both the temperature-sensitive fabA6 mutation and cold-sensitive secG null mutation of Escherichia coli. J Bacteriol 183(19): 5523–5528.

28. Baba, T., et al. (2006). Construction of Escherichia coli K-12 in-frame, single-gene knockout mutants: the Keio collection. Mol Syst Biol 2: 2006 0008.

29. Henry, M. F. and J. E. Cronan, Jr. (1991). Escherichia coli transcription factor that both activates fatty acid synthesis and represses fatty acid degradation. J Mol Biol 222(4): 843–849.

30. Campbell, J. W. and J. E. Cronan, Jr. (2001). Escherichia coli FadR positively regulates transcription of the fabB fatty acid biosynthetic gene. J Bacteriol 183(20): 5982–5990.

31. Nunn, W. D., et al. (1983). Role for fadR in unsaturated fatty acid biosynthesis in Escherichia coli. J Bacteriol 154(2): 554–560.

32. Pavoncello, V., et al. (2022). Degradation of Exogenous Fatty Acids in Escherichia coli. Biomolecules 12(8).

33. Cronan, J. E., Jr. and E. P. Gelmann (1973). An estimate of the minimum amount of unsaturated fatty acid required for growth of Escherichia coli. J Biol Chem 248(4): 1188–1195.

34. Heath, R. J., et al. (2001). Lipid biosynthesis as a target for antibacterial agents. Prog Lipid Res 40(6): 467–497.

35. Buttke, T. M. and L. O. Ingram (1978). Inhibition of unsaturated fatty acid synthesis in Escherichia coli by the antibiotic cerulenin. Biochemistry 17(24): 5282–5286.

36. Porrini, L., et al. (2013). Cerulenin inhibits unsaturated fatty acids synthesis in Bacillus subtilis by modifying the input signal of DesK thermosensor. Microbiologyopen 3(2): 213–24

37. Melchior, D.L. (1982) Lipid Phase Transitions and Regulation of Membrane Fluidity in Prokaryotes. Current Topics in Membranes and Transport 17, Pages 263-316

38. Marr, A. G. and J. L. Ingraham (1962). Effect of Temperature on the Composition of Fatty Acids in Escherichia Coli. J Bacteriol 84(6): 1260–1267.

39. 39. de Mendoza, D., and J. E. Cronan, Jr. (1983). Thermal regulation of membrane lipid fluidity in bacteria. Trends Biochem. Sci. 8: 49–52.

40. Gelmann, E.P. and J. E. Cronan, Jr. (1972). Mutant of *Escherichia coli* Deficient in the Synthesis of cis-Vaccenic Acid. J Bacteriol 112(1): 381–387.

41. 41. de Mendoza, D., et al. (1983). Thermal Regulation of Membrane Fluidity in *Escherichia coli*. J Biol Chem 258(4): 2098–2101.

42. Yao, J. and C. O. Rock (2017). Exogenous fatty acid metabolism in bacteria. Biochimie 141: 30–39.

43. Magnusson, L. U., et al. (2007). Identical, independent, and opposing roles of ppGpp and DksA in Escherichia coli. J Bacteriol 189(14): 5193–5202.

44. Anderson, S. E., et al. (2023). The transcription factor DksA exerts opposing effects on cell division depending on the presence of ppGpp. mBio 14(6): e0242523.

45. Du, S. and J. Lutkenhaus (2019). At the Heart of Bacterial Cytokinesis: The Z Ring. Trends Microbiol 27(9): 781–791.

46. Kanjee, U., et al. (2012). Direct binding targets of the stringent response alarmone (p)ppGpp. Mol Microbiol 85(6): 1029–1043.

47. Steinchen, W., et al. (2020). (p)ppGpp: Magic Modulators of Bacterial Physiology and Metabolism. Front. Microbiol.11:2072

48. Sanyal, R., et al. (2019). A Novel Gene Contributing to the Initiation of Fatty Acid Biosynthesis in Escherichia coli. J Bacteriol 201(19).

49. Sanyal, R. and R. Harinarayanan (2020). Activation of RelA by pppGpp as the basis for its differential toxicity over ppGpp in Escherichia coli. J Biosci 45.

50. Harinarayanan, R., et al. (2008). Synthetic growth phenotypes of Escherichia coli lacking ppGpp and transketolase A (tktA) are due to ppGpp-mediated transcriptional regulation of tktB. Mol Microbiol 69(4): 882–894.

51. Gohrbandt, M., et al. (2022). Low membrane fluidity triggers lipid phase separation and protein segregation in living bacteria. EMBO J 41(5): e109800.

52. Sinha, A.K., et al. (2019). Fatty acid starvation activates RelA depleting lysine precursor pyruvate. Mol Microbiol 112(4): 1339–1349.

53. Battesti, A. and E. Bouveret (2006). Acyl carrier protein/SpoT interaction, the switch linking SpoT-dependent stress response to fatty acid metabolism. Mol Microbiol 62(4): 1048–1063.

54. 54. Miller, J. (ed.). (1992). A short course in bacterial genetics: a laboratorymanual and handbook for Escherichia coli and related bacteria. Cold SpringHarbor Laboratory Press, Cold Spring Harbor, N.Y.

55. Sambrook J, Fritsch EF, Maniatis T. (1989). Molecular cloning: a laboratory manual, 2nd ed. Cold Spring Harbor Laboratory Press, Cold Spring Harbor, NY.

56. Baba, T., et al. (2006). Construction of Escherichia coli K-12 in-frame, single-gene knockout mutants: the Keio collection. Mol Syst Biol 2: 2006 0008.

57. Cherepanov, P. P. and W. Wackernagel (1995). Gene disruption in Escherichia coli: TcR and KmR cassettes with the option of Flp-catalyzed excision of the antibiotic-resistance determinant. Gene 158(1): 9–14.

58. Kitagawa, M., et al. (2005). Complete set of ORF clones of Escherichia coli ASKA library (a complete set of E. coli K-12 ORF archive): unique resources for biological research. DNA Res 12(5): 291–299.

59. Reddy, M. (2007). Role of FtsEX in cell division of Escherichia coli: viability of ftsEX mutants is dependent on functional SufI or high osmotic strength. J Bacteriol 189(1): 98–108.

60. Datsenko, K. A. and B. L. Wanner (2000). One-step inactivation of chromosomal genes in Escherichia coli K-12 using PCR products. Proc Natl Acad Sci U S A 97(12): 6640–6645.

61. Buyer, J. S. (2003). Improved fast gas chromatography for FAME analysis of bacteria. J Microbiol Methods 54(1): 117–120.

