## Supplementary Figures S1 to S7 for "(p)ppGpp buffers cell division when membrane fluidity decreases in *Escherichia coli*"

**Fig. S1**

LB Cm (30 °C) | LB Cm (42 °C)

+ 0.2% arabinose

*fabA2(ts)*/
pBAD33m

*fabA2*(*ts*)/
pBAD*fabA*

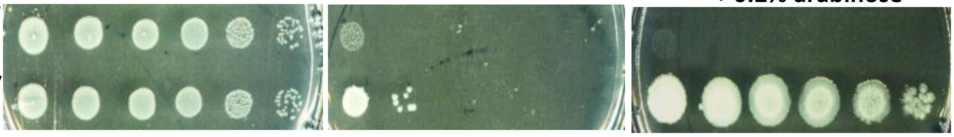

**Fig. S2**

ΔrelA ΔspoT Δ*fadR*/pRCspoT

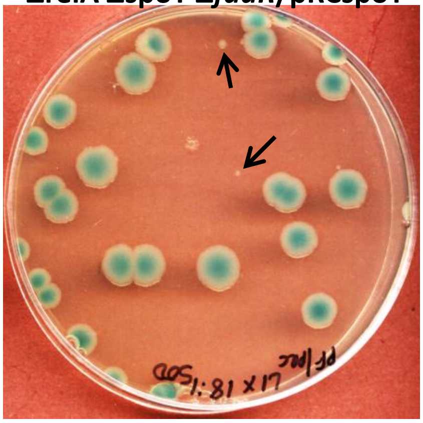

LB IPTG X-gal + 18:1 (30°C)

**Fig. S3**

Wild type

Δ*fabF*

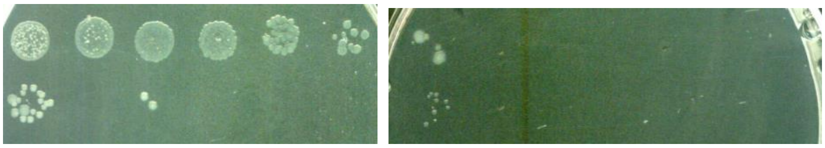

LB cer 40μg/ml | LB cer 80μg/ml

### Fig. S4

**A**

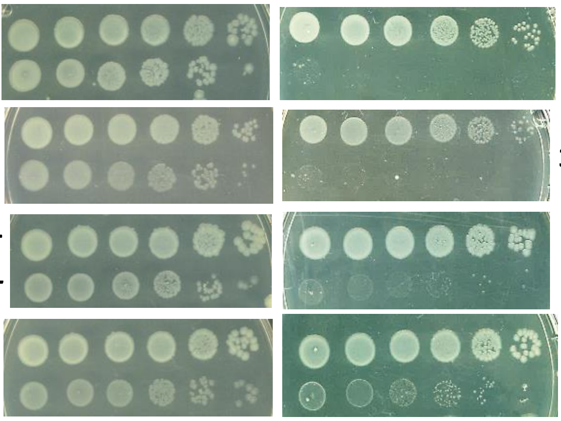


### Fig. S5

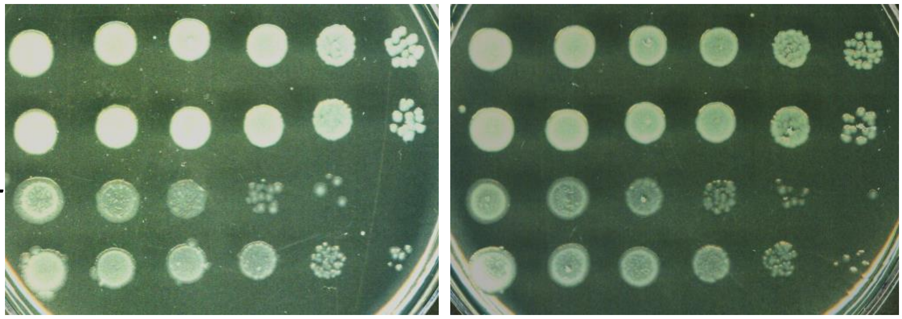

**Fig. S6**

**A**

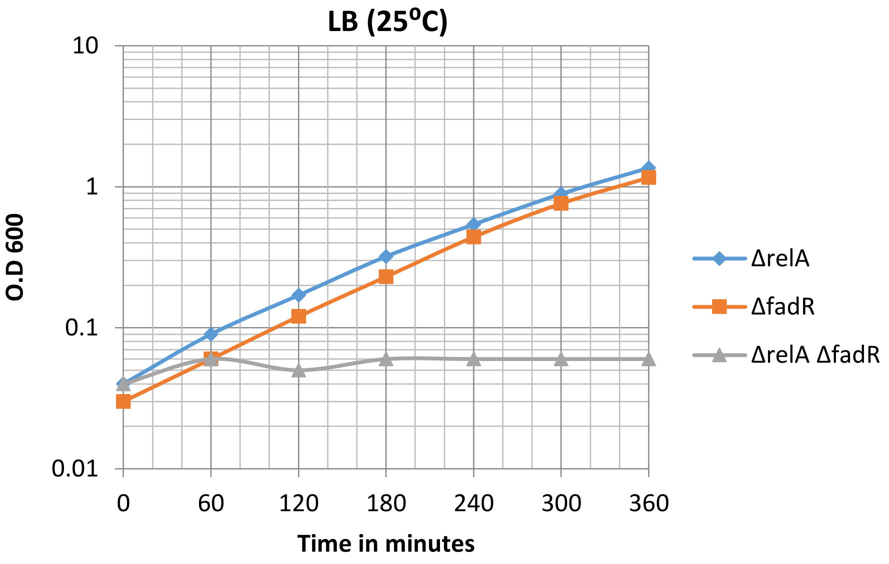
LB (25⁰C)
10
1
0.1
0.01
O.D 600
0
60
120
180
240
300
360
Time in minutes
ΔrelA
ΔfadR
ΔrelA ΔfadR

**B**

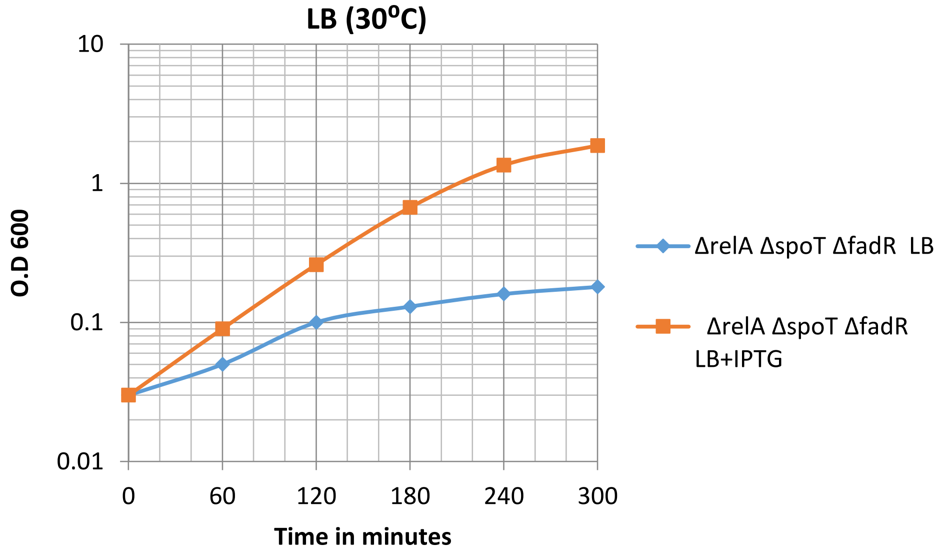
LB (30⁰C)
10
1
0.1
0.01
O.D 600
0
60
120
180
240
300
Time in minutes
ΔrelA ΔspoT ΔfadR LB
ΔrelA ΔspoT ΔfadR LB+IPTG

**Fig. S7**

(i)

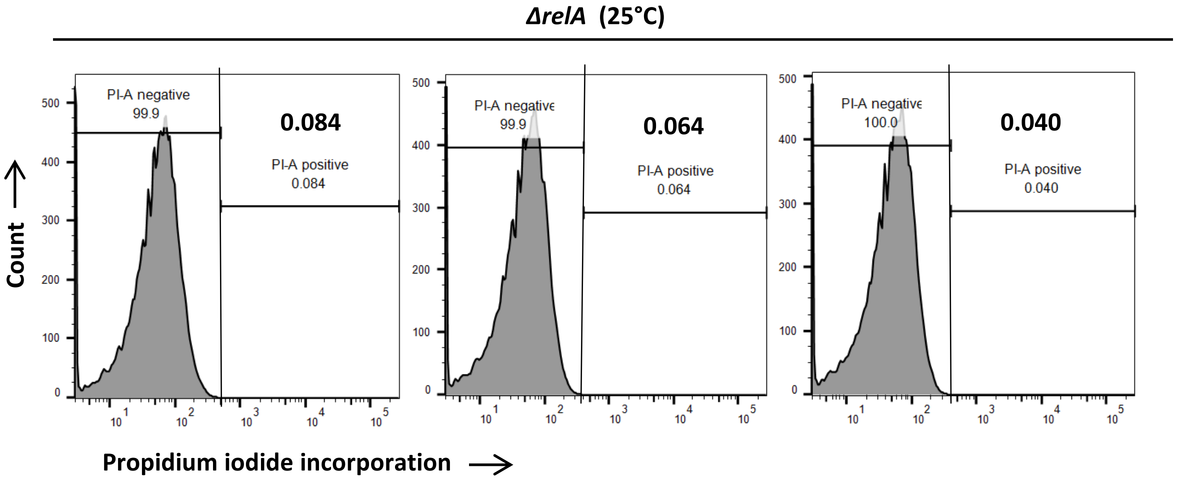


(ii)

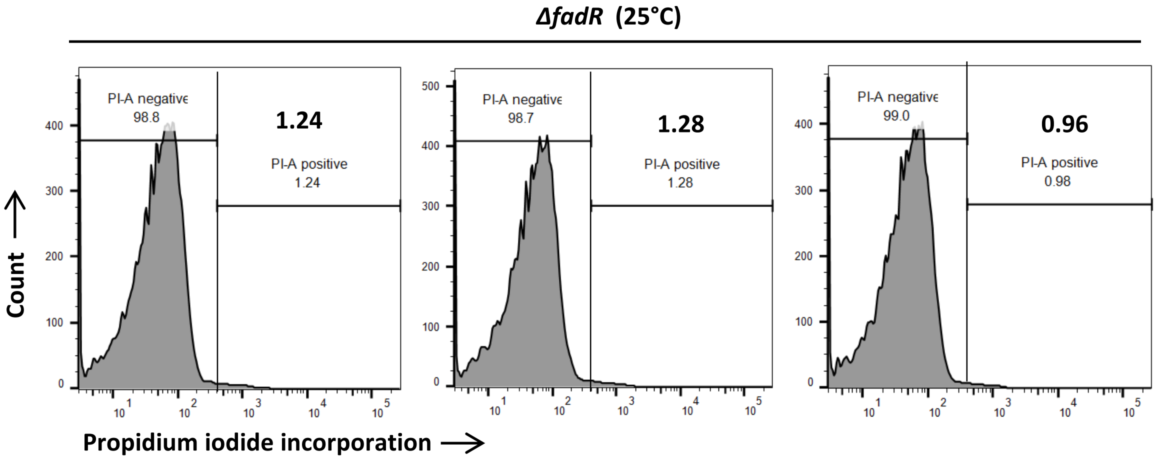


(iii)

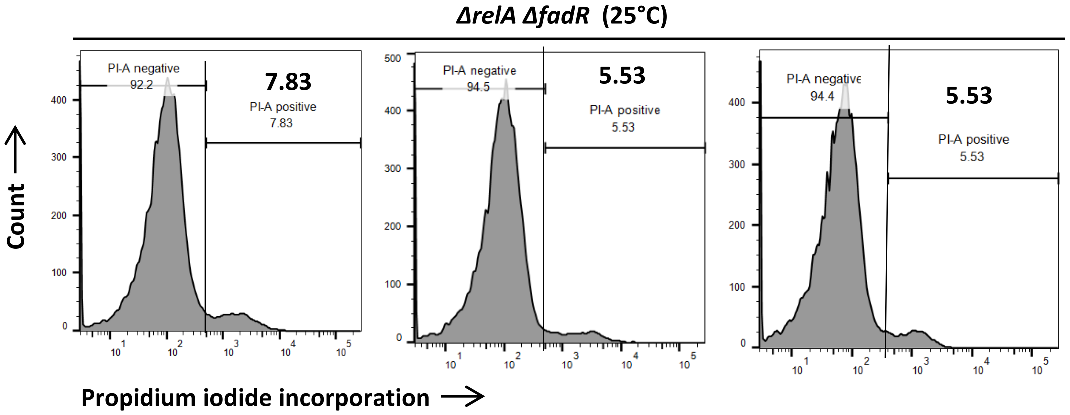

**Fig. S7 (continued)**

**(iv)** ***ΔrelA ΔspoT*** **(30°C)**

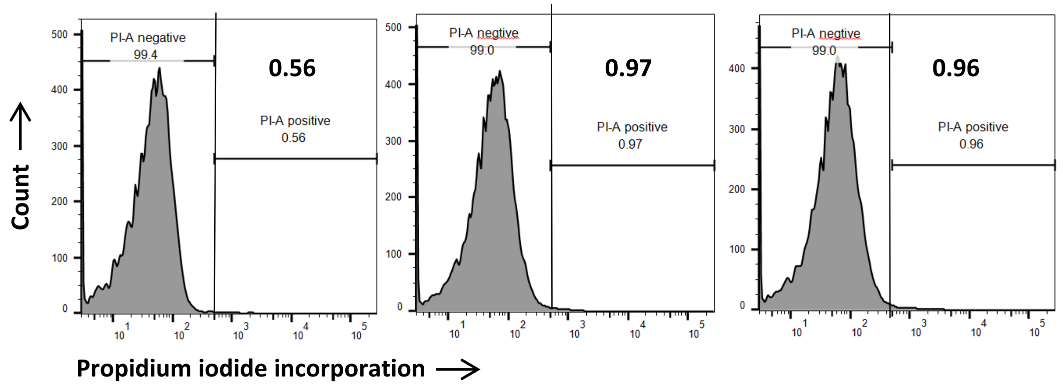


**(v)** ***ΔfadR*** **(30°C)**

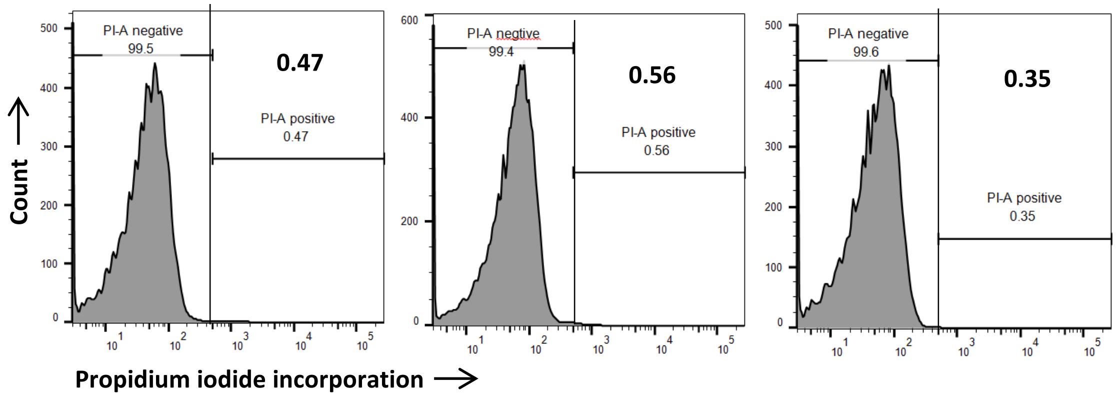


**(vi)** ***ΔrelA ΔspoT ΔfadR*****/pRC*****spoT*** **(30°C)**

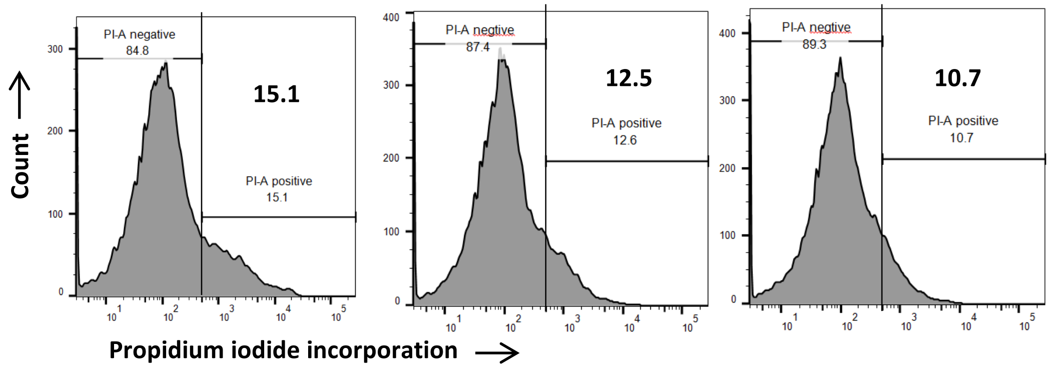
