## Supplementary Table 1 for "(p)ppGpp buffers cell division when membrane fluidity decreases in *Escherichia coli*"

**Table S1**

| Strains | | |
|---|---|---|
| **Strain** | **Genotype** | **Source/Reference** |
| MG1655 | F- $\lambda^-$ *ilvG*$^-$ *rfb-50 rph-1* (Wild type E.coli K-12) | Lab collection |
| CY50 | *F-, galK35, $\lambda^-$, fabA2(ts), trpC45, his-68, rpsL118(strR), malT1($\lambda^R$), xyl-7, mtlA2, thiE1* | CGSC, Yale. |
| CF10237 | CF1648 Δ*relA (relA256)* Δ*spoT (spoT212)* | Cashel lab |
| HR421 | MG1655 Δ*araBAD mal*::*lacI*$^q$ mini λtet | Lab collection |
| JW1176 | BW25113 Δ*fadR::Kan* (Keio collection) | (1) |
| JW2755 | BW25113 Δ*relA*::Kan (Keio collection) | (1) |
| JW1794 | BW25113 Δ*fadD*::Kan (Keio collection) | (1) |
| JW5020 | BW25113 Δ*fadE*::Kan (Keio collection) | (1) |
| JW1081 | BW25113 Δ*fabF*::Kan (Keio collection) | (1) |
| AN120 | Δ*relA* Δ*spoT* Δ*lacZYAI::FRT*/pRC*spoT* | (2) |
| VS2 | MG1655Δ*lacZYAI*::FRT | This work |
| VS3 | Δ*relA*::FRT Δ*lacZYAI*::FRT | This work |
| VS6 | Δ*relA* Δ*spoT* Δ*fadR*::Kan Δ*lacZYAI*::FRT /pRC*spoT* | This work |
| VS7 | Δ*relA* Δ*spoT* Δ*lacZYAI*::FRT Δ*fadR*::FRT/pRC*spoT* | This work |
| VS9 | Δ*fadR*::Kan Δ*lacZYAI*::FRT/pRC*spoT* | This work |
| VS11 | Δ*fadR*::FRT Δ*lacZYAI*::FRT | This work |
| VS14 | Δ*relA* Δ*spoT* Δ*lacZYAI*::FRT Δ*fadR*::Kan/pRC*spoT/* pCA*gnsB* | This work |
| VS15 | Δ*relA* Δ*spoT* Δ*fadR*::Kan Δ*lacZYAI*::Kan/pRC*spoT* /pCA24N | This work |
| VS16 | Δ*relA* Δ*spoT* Δ*fadR*::FRT Δ*lacZYAI*::FRT/pRC*spoT* /pCA24N | This work |
| VS22 | Δ*relA* Δ*spoT* Δ*fadR*::Kan Δ*lacZYAI*::FRT /pRC*spoT/* pCA*gnsA* | This work |
| VS23 | Δ*relA* Δ*spoT* Δ*fadR*::FRT Δ*lacZYAI*::FRT /pRC*spoT* /pCA*fabB* | This work |
| VS33 | Δ*relA*::FRT Δ*fadR*::Kan Δ*lacZYAI*::FRT | This work |
| VS75 | Δ*relA* Δ*spoT* Δ*fadR*::FRT Δ*fadD*::Kan Δ*lacZYAI*::FRT /pRC*spoT* | This work |
| VS76 | Δ*relA* Δ*spoT* Δ*fadR*::FRT Δ*fadE*::Kan Δ*lacZYAI*::FRT /pRC*spoT* | This work |
| VS119 | Δ*relA* Δ*spoT* Δ*fadR*::FRT Δ*lacZYAI*::FRT /pRC*spoT* /pBAD*fabA* | This work |
| VS120 | Δ*relA* Δ*spoT* Δ*fadR*::FRT Δ*lacZYAI*::FRT/pRC*spoT* /pBAD33m | This work |
| VS134 | Δ*relA*::FRT Δ*fadR*::Kan Δ*lacZYAI*::FRT/pCA*gnsB* | This work |
| VS135 | Δ*relA*::FRT Δ*fadR*::Kan Δ*lacZYAI*::FRT/pBAD33m | This work |
| VS136 | Δ*relA*::FRT Δ*fadR*::Kan Δ*lacZYAI*::FRT/pCA*fabB* | This work |
| VS137 | Δ*relA*::FRT Δ*fadR*::Kan Δ*lacZYAI*::FRT/pBAD*fabA* | This work |
| VS138 | Δ*relA*::FRT Δ*fadR*::Kan Δ*lacZYAI*::FRT/pCA*gnsA* | This work |
| VS149 | Δ*relA*::FRT Δ*fadR*::Kan Δ*lacZYAI*::FRT /pCA24N | This work |
| VS155 | MG1655 fabA2(ts) Δ*lacZYAI::FRT* | This work |
| VS157 | Δ*relA* Δ*spoT fabA2ts* Δ*lacZYAI::FRT* / pRC*spoT* | This work |

| | | |
|---|---|---|
| VS159 | Δ*relA*::FRT Δ*fadR*::FRT Δ*lacZYAI*::FRT | This work |
| VS210 | Δ*relA*::FRT Δ*fadR*::FRT Δ*fadE*::Kan Δ*lacZYAI*::FRT | This work |
| VS211 | Δ*relA*::FRT Δ*fadR*::FRT Δ*fadD*::Kan Δ*lacZYAI*::FRT | This work |
| VS418 | Δ*relA* Δ*spoT* Δ*fadD*::kan Δ*lacZYAI*::FRT /pRC*spoT* | This work |
| VS419 | Δ*relA* Δ*spoT* Δ*lacZYAI*::FRT Δ*fadE*::kan Δ*lacZYAI*::FRT /pRC*spoT* | This work |
| VS451 | Δ*relA* Δ*spoT* Δ*fadR*::FRT Δ*lacZYAI*::FRT/pRC*spoT* /pCL$_{Kan}$ 1920 | This work |
| VS452 | Δ*relA* Δ*spoT* Δ*fadR*::FRT Δ*lacZYAI*::FRT / pRC*spoT* / pCL$_{Kan}$ *ftsQAZ* | This work |
| VS469 | Δ*relA* Δ*spoT* Δ*lacZYAI*::FRT/pRC*spoT* /pCL$_{Kan}$1920 | This work |
| VS470 | Δ*relA* Δ*spoT* Δ*lacZYAI*::FRT/pRC*spoT* / pCL$_{Kan}$ *ftsQAZ* | This work |
| VS473 | MG1655 Δ*fabF*::Kan Δ*lacZYAI::*FRT | This work |
| VS474 | Δ*relA* Δ*spoT* Δ*fabF*::Kan Δ*lacZYAI*::FRT /pRC*spoT* | This work |
| VS478 | Δ*relA* Δ*spoT* Δ*lacZYAI*::FRT/pCA*fabB* | This work |
| VS479 | Δ*relA* Δ*spoT* Δ*lacZYAI*::FRT/pCA24N | This work |
| VS484 | MG1655 *fabA2ts* Δ*lacZYAI*::FRT/pBAD33m | This work |
| VS485 | MG1655 *fabA2tsΔlacZYAI::FRT*/pBAD*fabA* | This work |

| **Plasmids** | | |
|---|---|---|
| pRC7 | A low copy-number, mini-F derivative of pFZY1; the multiple cloning sites (MCS) in the *lac* promoter of pRC7 contain restriction sites for *Eco*RI, *Bam*HI, *Sal*I, and *Hin*dIII | (4) |
| pRCspoT | pRC7 which contains a minimal *spoT* ORF PCR amplified from MG1655 and ligated into *Eco*RI and *Hin*dIII sites, the expression of *spoT* is under *lac* promoter, therefore IPTG-inducible. | (2) |
| pRC$_{sp}$-spoT | The Sp$^{r}$ derivative made from pRC*spoT*, by replacing the *bla* gene with *aadA* by recombineering | (2) |
| pKD13 | Template plasmid with gene for kanamycin resistance flanked by FRT sites | (7) |
| pCA24N | Cloning vector (Cm$^{r}$) used in ASKA library construction. | (8) |
| pCAfabB | plasmid from ASKA plasmid collection which expresses *fabB* gene | (8) |
| pCAgnsA | plasmid from ASKA plasmid collection which expresses *gnsA* gene | (8) |
| pCAgnsB | plasmid from ASKA plasmid collection which expresses *gnsB* gene | (8) |

| | | |
|---|---|---|
| pBAD33 | An expression vector with a pACYC184 derived origin of replication and allows tightly regulated expression of the genes cloned under the PBAD promoter of the *araBAD* operon. In addition, the vector carries the *araC* gene, encoding the positive and negative regulator of this promoter. | (9) |
| pBAD33m | pBAD33m is a modified version of pBAD33 with *dhfR* gene flanked by *Nde*I and *Hin*dIII sites. | R. Varadarajan lab (IISc) |
| pBAD33mfabA | *fabA* gene with its native SD sequence was cloned in the pBAD33m vector between the *Kpn*I and *Hind*III sites | This work |
| pCL1920 | A pSC101-based, low copy number vector with spectinomycin (Sp) and streptomycin (Sr) resistance marker carrying the MCS in *lacZα* region and hence provides the advantage of screening the insertions using α-complementation | (10) |
| $pCL_{Kan}1920$ | The $Kan^r$ derivative made from pCL1920 , by replacing the *aadA* gene with *kanR* by Recombineering | This work |
| $pCL_{Kan}ftsQAZ$ | The $Kan^r$ derivative made from pCL*ftsQAZ*, by replacing the *aadA* gene with *kanR* by recombineering | This work |

| **Primers** | |
|---|---|
| JGABkpnfabAFP | ATATATATATATGGTACCTTCAATAAAATAAGGCTTACAGAGAAC |
| JGABfabAH3RP | ATATATATATATAAGCTTTCAGAAGGCAGACGTATCCTGGAAC |
| JGABKanpKD13FP | CGCGAAGCGGCGTCGGCTTGAACGAATTGTTAGACATTAGAAGAA CTCGTCAAGAAGGCG |
| JGABKanpKD13RP | CAGCAGGGCAGTCGCCCTAAAACAAAGTTAAACATCATGATTGAA CAAGATGGATTGCACG |

**Reference**


1. Baba, T., et al. (2006). Construction of *Escherichia coli* K-12 in-frame, single-gene knockout mutants: the Keio collection. Mol Syst Biol **2**: 2006 0008.

2. Nazir, A. and R. Harinarayanan (2015). Inactivation of Cell Division Protein FtsZ by SulA Makes Lon Indispensable for the Viability of a ppGpp0 Strain of Escherichia coli. J Bacteriol **198**(4): 688-700.

4. Bernhardt, T. G. and P. A. de Boer (2004). Screening for synthetic lethal mutants in Escherichia coli and identification of EnvC (YibP) as a periplasmic septal ring factor with murein hydrolase activity. Mol Microbiol **52**(5): 1255-1269.

7. Datsenko, K. A. and B. L. Wanner (2000). One-step inactivation of chromosomal genes in Escherichia coli K-12 using PCR products. Proc Natl Acad Sci U S A **97**(12): 6640-6645.

8. Kitagawa, M., et al. (2005). Complete set of ORF clones of Escherichia coli ASKA library (a complete set of E. coli K-12 ORF archive): unique resources for biological research. DNA Res **12**(5): 291-299.

9. Guzman, L. M., et al. (1995). "Tight regulation, modulation, and high-level expression by vectors containing the arabinose PBAD promoter." J Bacteriol **177**(14): 4121-4130.

10. Lerner, C. G. and M. Inouye (1990). Low copy number plasmids for regulated low-level expression of cloned genes in Escherichia coli with blue/white insert screening capability. Nucleic Acids Res **18**(15): 4631.
